# Gut miRNA regulates gut microbiome and Alzheimer’s pathology in *App*-knock-in mice

**DOI:** 10.1101/2025.06.05.658056

**Authors:** Wenlin Hao, Qinghua Luo, Ilona Magdalena Szabo, Gilles Gasparoni, Sascha Tierling, Wenqiang Quan, Hsin-Fang Chang, Gang Wu, Julia Schulze-Hentrich, Yang Liu

## Abstract

Alzheimer’s disease (AD) alters the gut microbiome. It remains unclear whether manipulation of miRNA could modulate the gut microbiome and AD pathologies. Depletion of gut miRNA in *App*-knock-in mice by conditional knockout of *Dicer1* gene in intestinal epithelial cells: 1) decreased the absolute number and changed the composition of bacteria in both the gut and brain; 2) reduced cerebral Aβ load by inhibiting β-secretase activity and increasing the expression of LRP1 and ABCB1 at the blood-brain barrier; 3) increased *Il-10* transcription and decreased the transcription of *Ccl-2* gene, and that of *Ndufa2* and *Ndufa5* genes encoding mitochondrial respiratory enzymes in the brain; and 4) induced anxiety symptoms without affecting cognitive function in AD mice. Thus, manipulating miRNA in the gut can modify AD pathogenesis. Future studies should focus on identifying AD-specific miRNAs in the gut that can be therapeutically exploited (e.g. by oral administration) to prevent the progression of AD.

## Introduction

The mammalian gut contains an extraordinary number of commensal bacteria that influence the brain directly via the vagus nerve and indirectly through the release of metabolites and neurotransmitters as well as by educating immune cells [1, 2]. In patients of Alzheimer’s disease (AD), the composition of gut bacteria changes at the preclinical stage [3, 4], correlating with the inflammatory state of peripheral blood circulation [5], and the severity of cognitive deficits and brain pathology [3, 6, 7]. Sporadic use of antibiotics was reported to potentially decrease the risk of dementia in older adults after adjustment for comorbidities [8]. Transplantation of intestinal bacteria from AD patients impaired neurogenesis and cognitive function in gut bacteria-depleted rats [9]. In Alzheimer’s precursor protein (APP)-transgenic or knock-in mice, depletion of gut bacteria inhibits microglial inflammatory responses to amyloid-β peptide (Aβ) [10–13], and reduces Aβ deposits in the brain [13–19]. All these findings support the intricate relationship between gut microbiota and AD.

However, AD-associated profile of gut microbiome has not been defined, although bacteria from *Firmicutes* phylum often decrease, while those from *Proteobacteria* phylum increase [3, 5, 20]. The abundance of *Bacteroidetes* bacteria either decreases [6, 20, 21] or increases [20, 22] in various studies. Many *Firmicutes* bacteria produce butyrate, maintain intestinal barrier integrity, avoid excessive intestinal inflammation and prevent the progression of AD [5, 23–26]; in contrast, *Proteobacteria* bacteria, including from *Escherichia/Shigella* genus, release toxins and trigger inflammatory activation [5]. At the family and genus level, the results on gut microbiome in AD patients vary greatly from study to study; for example, a consistent decrease in butyrate-producing bacteria of the genus *Butyricicoccus* or *Coprococcus* or an increase in the genera *Escherichia/Shigella* (three genera are not changed in the same study) in AD patients is only found in two out of twenty independent studies [25]. The variable changes in gut microbiome also exist in different APP-transgenic AD mice [25]. Therefore, a potential precision therapy for each AD patient that targets potential AD-associated bacterial taxa in the gut should be considered. Our recent study suggests that depletion of a specific group of gut bacteria that stimulate interleukin (IL)-17a-expressing T lymphocytes may be able to prevent AD progression in APP-transgenic mice [13]. A flexible method to efficiently and specifically modify the microbiome in AD patients and animal models is desirable.

Intestinal epithelial cells have been shown to release microRNA (miRNA), a small non-coding RNA molecule, which enters intestinal bacteria, post-transcriptionally modifies bacterial protein expression and subsequently alters bacterial proliferation and metabolism [27]. While we found no literature directly documenting the association between microbiome and miRNA in the gut of AD patients or animals, a recent study has demonstrated this association in pediatric patients with active Crohn’s disease [28]. Interestingly, oral administration of miR-30d that is enriched in the feces of untreated multiple sclerosis patients and mice with experimental autoimmune encephalomyelitis (EAE) at the peak disease was reported to attenuate symptoms of EAE mice by increasing *Akkermansia muciniphila* bacteria and the subsequent population of regulatory T cells in the gut [29]. Before we determine the specific changes in gut microbiome and miRNA in AD mice and patients, and to feed AD mice with a specific miRNA for a long term (≥ 2 months), we knocked out *Dicer1* gene that encodes the key enzyme for miRNA production in intestinal epithelial cells and investigated the effects of overall miRNA depletion on AD-associated brain pathology in *App^NL-G-F^* knock-in mice.

## Materials and Methods

### Animal models and cross-breeding

*App^NL-G-F^* knock-in (App^ki/ki^) mice were kindly provided by T.C. Saido, RIKEN Brain Science Institute, Japan [30], *Dicer1*-floxed (Dicer1^fl/fl^) mice by A. Domanskyi, University of Helsinki, Finland [31] and Villin-Cre mice over-expressing CreERT2 under control of the *Villin* gene promoter by E. Batlle, The Barcelona Institute of Science and Technology (BIST), Spain [32]. App^ki/ki^ mice express humanized *APP* in the endogenous *App* locus harboring Swedish, Beyreuther/Iberian and Arctic mutations, avoiding the transgenic mouse artifacts that Aβ is generated along with overproduction of other APP fragments. The mice show typical Aβ pathology, neuroinflammation and memory impairment in an age-dependent manner [30]. App^ki/ki^ mice with and without expression of dicer1 in epithelial cells of the intestine (App^ki/ki^Dicer1^del^ and App^ki/ki^Dicer1^wt^) were created by crossing App^ki/ki^, Dicer1^fl/fl^ and Villin-Cre mice to obtain the genotypes App^ki/ki^/Dicer1^fl/fl^/Cre^tg^ and App^ki/ki^/Dicer1^fl/fl^/Cre^wt^, followed by feeding littermate AD mice with 600mg/kg tamoxifen (Carbolution Chemicals GmbH, St. Ingbert, Germany) supplements in diet from 6 month of age for 1 month. AD phenotypes were analyzed at 9 months before the end of the experiments. During experiments, the experimenters did not know which genotype the mice had.

All App^ki/ki^Dicer1^del^ and App^ki/ki^Dicer1^wt^ mice on a C57BL/6 genetic background born between June 2018 and December 2019 were included in this study. As we only compared AD phenotypes between dicer1-deficient and wild-type brothers or sisters, all mouse litters without suitable comparison pairs were excluded. Both male and female mice were used, with the proportion of male mice generally being 41.49%. The experimental littermate mice with different expression of CreERT2 were kept in the same cage and received tamoxifen-supplemented diet. To exclude a sex-related bias in AD pathology, we performed a preliminary experiment to compare Aβ42 levels in brain homogenates from 9-month-old male and female App^ki/ki^ mice. We found no significant effect of sex on cerebral Aβ level (see Supplementary Figure 1). Animal experiments were conducted in accordance with national rules and ARRIVE guidelines, and authorized by Landesamt für Verbraucherschutz, Saarland, Germany (registration numbers: 36/2018).

### Behavior tests

An open field test and a Y-maze test were performed (see the setting presented in Fig. 8, A and F). In the open field test, each mouse was allowed to run freely for 10 minutes in a bare and open chamber. Initially, the mice often stayed close to the walls, and later explored the center area. The total distance, duration in the center area, frequency of visits to the center area and velocity were monitored with a camera placed above the open field. The open field test was conducted once a day for three consecutive days. The Y-maze was made of gray plastic and consisted of three compartments radiating out from the center platform. In this test, each mouse was placed in the center of the maze facing toward one of the arms and was then allowed to explore freely for 5 minutes. The number of arm entries and the number of triads were recorded to calculate the percentage of alternation. An entry occurred when all four limbs were within the arm. The Y-maze test was repeated three times per day in three consecutive days.

### Tissue collection and isolation of blood vessels

Animals were euthanized by inhalation of overdose isoflurane. After intracardial perfusion with ice-cold phosphate-buffered saline (PBS), the brain was removed and divided along the anterior-posterior axis. The left hemisphere was immediately fixed in 4% paraformaldehyde (Sigma-Aldrich Chemie GmbH, Taufkirchen, Germany) in PBS and embedded in paraffin for immunohistochemistry. The cortex and hippocampus were carefully separated from the right hemisphere. A roughly 0.5-mm thick sagittal section of tissue was cut from the medial side and homogenized in TRIzol (Thermo Fisher Scientific, Darmstadt, Germany) for RNA isolation. The remainder was snap-frozen in liquid nitrogen and stored at - 80°C until biochemical analysis. The cecum together with a 0.5-cm-long segment of the neighboring colon was also collected, snap-frozen and stored at -80°C for isolation of intestinal bacteria.

To isolate brain blood vessels, the cortex and hippocampus from 9-month-old App^ki/ki^Dicer1^del^ and App^ki/ki^Dicer1^wt^ mice were dissected and brain vessel fragments were isolated as we did previously [33]. Briefly, brain tissues were homogenized in HEPES-contained Hanks’ balanced salt solution (HBSS) and centrifuged at 4,400g in HEPES-HBSS buffer supplemented with dextran from *Leuconostoc spp*. (molecular weight ∼70,000; Sigma-Aldrich) to delete myelin. The vessel pellet was re-suspended in HEPES-HBSS buffer supplemented with 1% bovine serum albumin (Sigma-Aldrich) and filtered with 20 μm-mesh. The blood vessel fragments were collected on the top of filter and stored at -80°C for biochemical analysis.

### 16S rDNA sequencing and microbiome analysis in the gut and brain

Bacterial DNA was extracted from intestinal bacteria (100mg) in the frozen cecum and neighboring colon with QIAamp Fast DNA Stool Mini Kit (Qiagen, Hilden, Germany). The amount of bacteria was evaluated by quantifying 16S rDNA with SYBR Green-based real-time PCR using the universal bacterial r16S gene primers (16S-V2-101F: 5-AGYGGCGIACGGGTGAGTAA-3, and 16S-V2-361R: 5-CYIACTGCTGCCTCCCGTAG-3) as we did in a previous study [13]. The amount *Gapdh* DNA derived from the intestinal cells was also measured and used as an internal control with the primers: sense, 5’-*ACAACTTTGGCATTGTGGAA*-3’ and antisense, 5’-*GATGCAGGGATGATGTTCTG*-3’. The V3 - V4 region of the 16S rRNA-encoding gene was then amplified with the barcode fusion primers (338F and 806R). After purification, PCR products were used for constructing libraries and sequenced on the Illumina MiSeq platform at Majorbio Co. Ltd. (Shanghai, China). The raw data was processed on Qiime2 (https://qiime2.org/), reducing sequencing and PCR errors, and denoising to get the operational taxonomic unit (OTU) consensus sequences, which were mapped to the 16S Mothur-Silva SEED r119 database (http://www.mothur.org/). Alpha diversity including Sobs, Shannon, Ace, Chao and Simpson indexes were used for the analysis of bacterial richness and diversity in a single mouse. Principal coordinate analysis (PCoA) and analysis of similarity (ANOSIM) were used for β-diversity analysis to compare bacterial compositions on genus level between App^ki/ki^Dicer1^del^ and App^ki/ki^Dicer1^wt^ mice. The difference of bacterial compositions on genus level between these two groups were also compared with Wilcoxon rink sum test. Additionally, we employed the Phylogenetic Investigation of Communities by Reconstruction of Unobserved States (PICRUSt2) software [34] to predict microbial functions by annotating the gene catalog based on the Kyoto Encyclopedia of Genes and Genomes (KEGG) modules [35]. All the analysis was performed using cloud-based tools with default analysis parameters (https://cloud.majorbio.com/page/tools.html).

Total DNA was isolated from mouse brains using TRIzol. Real-time PCR was used as described above to quantify the bacterial 16S rDNA in the brain tissue with murine *Gapdh* DNA as an internal control. Subsequently, the V3-V4 region of bacterial 16S rDNA was sequenced and analyzed for the microbiome at Novogene GmbH, Munich, Germany. Aamplicon sequence variants (ASVs) instead of OUT were generated from the denoised raw data, and annotated with the database Silva 138.1. The α- and β-diversity analysis were performed. The difference in bacterial abundance in the brains of App^ki/ki^Dicer1^del^ and App^ki/ki^Dicer1^wt^ mice at the genus level was determined by MetagenomeSeq analysis [36].

### Positive selection of CD4-positive cells from the spleen

The spleen of 9-month-old AD mice were prepared for single-cell suspensions using Spleen Dissociation Kit (mouse) (Miltenyi Biotec GmbH, Bergisch Gladbach, Germany). After blocking with 50 µg/ml CD16/CD32 antibody (clone 2.4G2; BioXCell, Lebanon, USA), CD4-positive spleen cells from the spleen with Dynabeads magnetic beads-conjugated CD4 antibody (clone L3T4; Thermo Fisher Scientific). Lysis buffer was immediately added to selected cells and total RNA was isolated using RNeasy Plus Mini Kit (Qiagen).

### Histological analysis

Serial 50-μm-thick sagittal sections were cut from the paraffin-embedded hemisphere. Four neighboring sections with 300 µm of interval were deparaffinized, labeled with rabbit anti-ionized calcium-binding adapter molecule (Iba)-1 antibody (Wako Chemicals, Neuss, Germany) and VectaStain *Elite* ABC-HRP kit (Cat.-No.: PK-6100, Vector Laboratories, Burlingame, USA), and visualized with diaminobenzidine (Sigma-Aldrich). Iba-1-positive microglia/brain macrophages were counted with Optical Fractionator in the hippocampus on a Zeiss AxioImager.Z2 microscope (Carl Zeiss Microscopy GmbH, Göttingen, Germany) equipped with a Stereo Investigator system (MBF Bioscience, Williston), as we did previously [37].

To evaluate the cerebral Aβ level, after deparaffinization, 4 serial brain sections from each animal were stained with mouse monoclonal Aβ antibody (clone MOAB-2; BioXCell, Lebanon, USA), biotin-conjugated goat anti-mouse IgG and Cy3-conjuagted streptavidin (Jackson ImmunoResearch Europe Ltd. Cambridge, UK). After mounting, the whole section including hippocampus and cortex was imaged with Microlucida (MBF Bioscience) and merged. The positive staining and brain region analyzed were measured for the area with Image J tool “Analyse Particles” (https://imagej.nih.gov/ij/). The threshold for all compared samples was manually set and kept constantly. The percentage of Aβ coverage in the brain was calculated.

### Western blot analysis

Frozen brain tissues and blood vessel isolates were homogenized in RIPA buffer (50mM Tris [pH 8.0], 150mM NaCl, 0.1% SDS, 0.5% sodiumdeoxy-cholate, 1% NP-40, and 5mM EDTA) supplemented with protease inhibitor cocktail (Roche Applied Science, Mannheim, Germany) on ice. The proteins were separated by 10% or 12% Tris-glycine SDS/PAGE. Before loading on PAGE gel, vessel preparations were sonicated. Western blots were performed using rabbit monoclonal antibodies against ABCB1 (clone E1Y7S), PDGFRβ (clone 28E1), CD13/APN (clone D6V1W), Munc18-1 (clone D4O6V), SNAP25 (clone D110), PSD95 (clone D74D3), and synaptophysin (clone D8F6H), and rabbit polyclonal antibody against LRP1 (Cat.-No.: 64099) (all antibodies above were bought from Cell Signaling Technology), as well as rabbit polyclonal antibody against claudin-5 (Cat.-No.: GTX49370; GeneTex, Hsinchu, China). The detected proteins were visualized via the Plus-ECL method (PerkinElmer, Waltham, USA). To quantify proteins of interest, rabbit monoclonal antibody against β-actin (clone 13E5) or Vinculin (clone E1E9V), or rabbit polyclonal antibody against α-tubulin (Cat.-No.: 2144) (these three antibodies were from Cell Signaling Technology) was used as a protein loading control. Densitometric analysis of band densities was performed with Image-Pro Plus software version 6.0.0.260 (Media Cybernetics, Rockville, MD). For each sample, the protein level was calculated as a ratio of target protein/loading control per sample.

### Brain homogenates and ELISA assays of Aβ

The frozen brain hemispheres were homogenized and extracted serially in Tris-buffered saline (TBS), TBS plus 1% Triton X-100 (TBS-T), guanidine buffer (5 M guanidine HCl/50 mM Tris, pH 8.0) as described in our previous study [37]. Aβ concentrations in three separate fractions of brain homogenates were determined by Human Amyloid β42 and β40 Brain ELISA kits (Cat.-No.: EZBRAIN42 and EZBRAIN40, respectively; both from Merck Chemicals GmbH, Darmstadt, Germany). Results were normalized on the basis of the sample’s protein concentration.

### mRNA seq library preparation, sequencing, processing and analysis

Total RNA was isolated from mouse brains using TRIzol. RNAseq libraries were prepared using 100 ng of total RNA per sample with the NEBnext Ultra II Directional RNA library kit for Illumina (New England Biolabs [NEB]; Ipswich, USA) and the NEBNext Poly(A) mRNA Magnetic Isolation Module (NEB) according to the manufacturer’s recommendations. Libraries were quantified with the NEBnext library quantification kit for Illumina (NEB) and then sequenced on an Aviti sequencing system (Element Biosciences; San Diego, USA) for 30 – 40 million 2× 75 nt reads per sample. Raw data was processed with the nf-core RNAseq pipeline version 3.17.0 (https://nf-co.re/rnaseq/3.17.0/) and nextflow v23.04.1 applying the default settings and using mm10 reference assembly. Differential analysis was performed in R using the DESeq2 package. Gene ontology (GO) enrichment analysis was performed using the GO Consortium-provided website-based free service (https://geneontology.org) with a 0.05 adjusted *p*-value cut-off and a minimum absolute log2fold change of 1, and also using the GOstats package (https://www.bioconductor.org/packages/release/bioc/html/GOstats.html) with a 0.05 *p*-value cut-off and a minimum absolute log2fold change of 1.

### Quantitative PCR for analysis of gene transcripts

Total RNA was isolated from mouse brains or the wall of cecum with TRIzol or from selected CD4-positive spleen cells with RNeasy Plus Mini Kit (Qiagen) and reverse-transcribed. Gene transcripts were quantified with established protocols [37, 38] and Taqman gene expression assays of mouse *Tnf-α*, *Il-1β*, *Chemokine (C–C motif) ligand 2* (*Ccl-2*), *Il-10*, *Chitinase-like 3* (*Chi3l3*), *Allograft inflammatory factor 1* (*AIF-1*, encoding Iba-1 protein), *Il-17a*, *RAR-related orphan receptor C* (*Rorc*), *Interferon-γ* (*Ifn-γ*), *Il-4*, *Neprilysin* and *Insulin-degrading enzyme* (*Ide*), *Gapdh* and 18S ribosomal RNA (*18S*) (Thermo Fisher Scientific). The transcription of *mt-Nd4, mt-Nd4l, mt-Atp6, Atp5e*, *Atp6v1c2*, *Ndufa2*, *Ndufa5*, *Romo1*, *Lrrc17*, *Agrp*, *Zo1*, *Occludin1*, *Claudin1*, *Claudin2*, *Claudin5* and *Dicer1* genes in brain tissues was determined using the SYBR green binding technique with the primers showed in Table 1, part of which were copied from the published studies [39, 40].

**Table 1.**
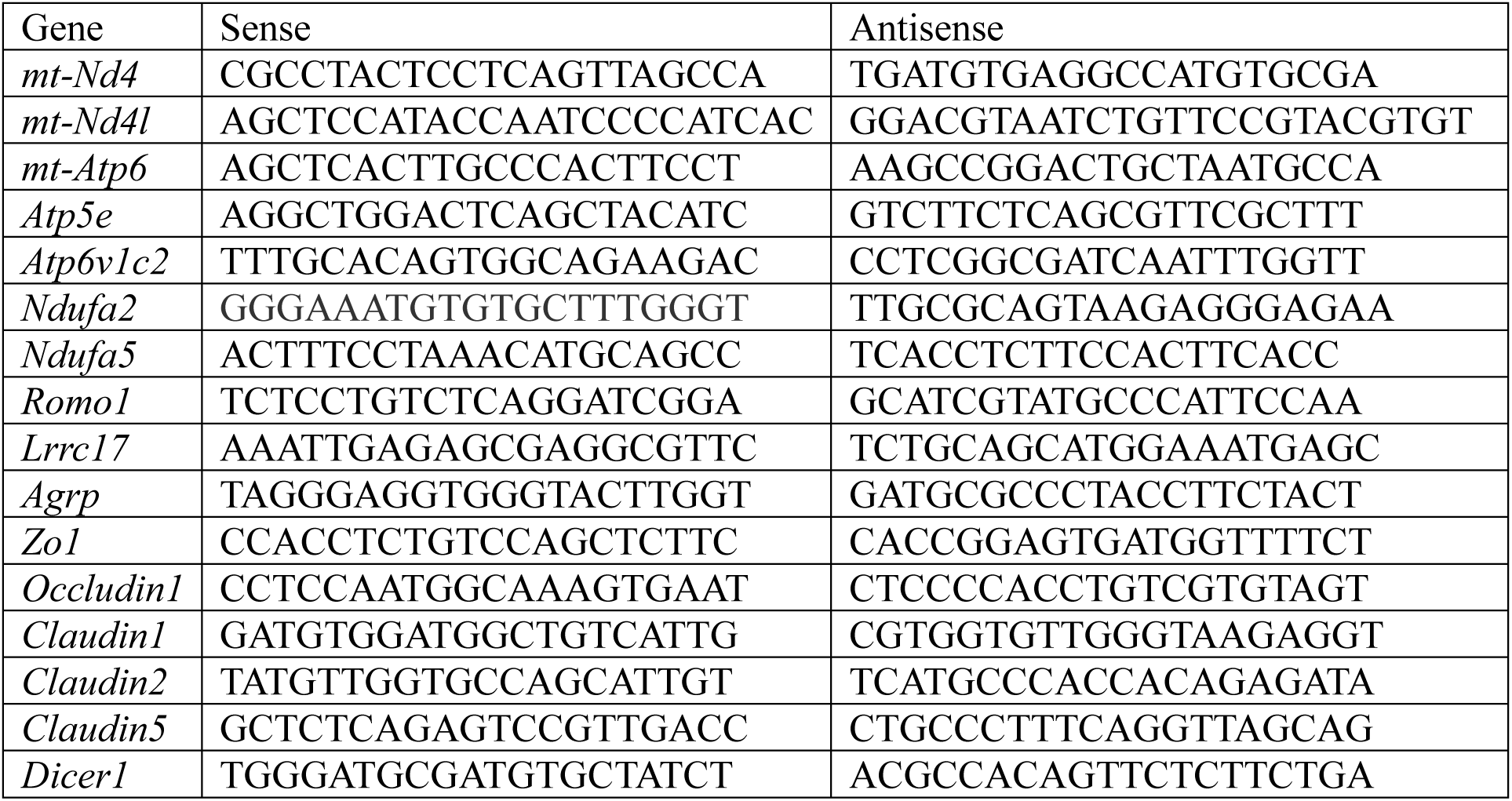
List of primer sequences used for SYBR green-based quantitative RT-PCR.

### Statistical analysis

Data were presented as mean ± *SEM* and displayed using a scatter plot with a bar overlay in the figure, with the scatter plot representing individual data points. Continuous variables between two groups of cases were compared with two independent-samples Students *t*-test or Mann-Whitney-*U*-test, depending on whether the variables were normally distributed. To exclude the possibility that the reduction of cerebral Aβ was co-caused by the depletion of miRNA in the gut and sex, we performed a linear regression analysis with Aβ as the dependent variable and the deletion of epithelial Dicer1 and sex as independent variables. For multiple comparisons, we used one-way or two-way ANOVA followed by Bonferroni, Tukey, or Dunnett T3 *post hoc* test (dependent on the result of Levene’s test to determine the equality of variances). All statistical analyses were performed with GraphPad Prism 8 version 8.0.2. for Windows (GraphPad Software, San Diego, USA) or SPSS software for Windows (Version 26.0, IBM, Armonk, USA). Other statistical methods in microbiome analysis offered by external companies have been described above. Statistical significance was set at *p* < 0.05.

## Results

### Establishment of *App*-knock-in mice with deletion of dicer1 in gut epithelial cells

Intestinal epithelial cells are the major source of miRNA in the intestinal lumen [27]. Six-month-old App^ki/ki^Dicer1^del^ and App^ki/ki^Dicer1^wt^ mice were administered with oral tamoxifen for one month to delete dicer1 in the intestinal epithelial cells, and analyzed for the phenotype in the gut at 9 months of age. Due to technical difficulties, we were unable to directly measure miRNA amount in the lumen of gut. However, we observed that the ratio of *Dicer1/18S* gene transcripts in the cecum wall was significantly lower in App^ki/ki^Dicer1^del^ mice than in App^ki/ki^Dicer1^wt^ mice (Fig. 1, A; 4.522 ± 0.851 × 10^-6^ vs. 9.167 ± 1.776 × 10^-6^; *t* test, *p* < 0.05). We also observed a significant reduction in transcription of *Claudin 1* gene but no other tight junction protein (TJP)-encoding genes in the cecum (Fig. 1, B - F; *t* test, *p* < 0.05). Our results were consistent with previous studies on the gut miRNA [27, 32], suggesting successful deletion of dicer1 in the gut epithelial cells of our AD mice.

**Figure 1.**
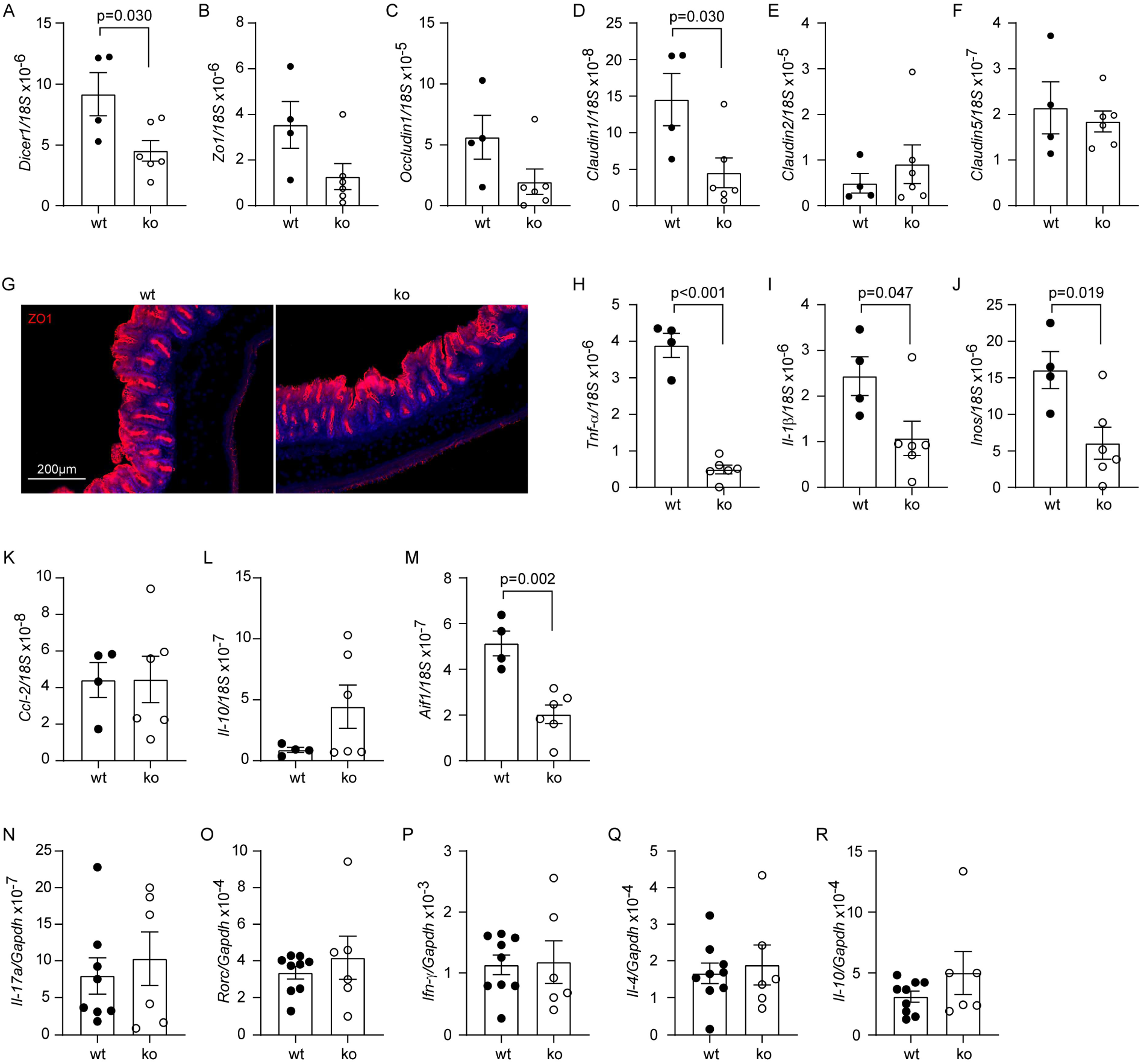
Deletion of dicer1 in gut epithelial cells did not impair the integrity of intestinal barrier in *App*-knock-in mice. Nine-month-old *App*-knock-in mice with (ko, n = 6) and without (wt, n = 4) deletion of dicer1 in gut epithelial cells were detected for gene transcription in the wall of cecum and imaged for the cecum morphology (A - M). Deletion of dicer1 significantly reduced the transcription of *Dicer1* and *Claudin1* genes (A and D; *t* test), as well as the transcription of various inflammatory genes, *Tnf-α*, *Il-1β*, *Inos* and *Aif1* (H - J and M; *t* test), but did not change the morphology of gut (G). Total RNA was also extracted from CD4-positive splenocytes of dicer1- deleted and wild-type *App*-knock-in mice and measured for mRNA levels using quantitative RT-PCR. Deletion of dicer1 did not alter the transcription of *Il-17a*, *Rorc*, *Ifn-γ*, *Il-4* and *Il-10* genes (N - R; *t* test, *n* = 6 - 9 per group).

Reduced TJP expression may impair the intestinal barrier, therefore, we examined the structure and inflammatory activation of the gut. Intestinal morphology after ZO1 staining did not differ between App^ki/ki^Dicer1^del^ and App^ki/ki^Dicer1^wt^ mice (Fig. 1, G). The transcription level of inflammatory genes, e.g., *Tnf-α*, *Il-1β*, *Inos*, and *Aif-1* that encodes Iba-1 protein, but not that of *Ccl-2* and *Il-10* genes, was significantly lower in App^ki/ki^Dicer1^del^ mice than in App^ki/ki^Dicer1^wt^ mice (Fig. 1, H - M; ; *t* test, *p* < 0.05). Moreover, deletion of *Dicer1* gene in the gut did not alter gene transcription of *Il-17a*, *Rorc*, *Ifn-γ*, *Il-4* and *Il-10* in CD4-positive splenocytes of App^ki/ki^Dicer1^del^ mice compared to App^ki/ki^Dicer1^wt^ mice (Fig. 1, N - R; *t* test, *p* > 0.05). All these results suggest that deletion of dicer1 in epithelial cells does not result in damage to intestinal barrier integrity, as leakiness of the gut barrier should trigger inflammatory activation and immune response in the gut. Our dicer1-deleted App^ki/ki^ mice allowed us to investigate the effects of miRNA on bacteria in the gut and subsequent AD pathologies in the brain.

### Deletion of dicer1 in gut epithelial cells modifies gut microbiome in *App*-knock-in mice

We knocked out *Dicer1* gene specifically in epithelial cells in App^ki/ki^Dicer1^del^ mice, which should reduce miRNA in the gut compared with App^ki/ki^Dicer1^wt^ mice. Indeed, 2 months after tamoxifen induction, the number of bacteria as shown by the level of bacterial 16S rDNA was significantly reduced in cecum contents of 9-month-old App^ki/ki^Dicer1^del^ mice compared with App^ki/ki^Dicer1^wt^ littermates (Fig. 2, A; *t* test, *p* < 0.05). In α-diversity analysis of bacterial species, Sobs, Ace, Chao, and Shannon indexes were significantly smaller in App^ki/ki^Dicer1^del^ mice than in App^ki/ki^Dicer1^wt^ littermates (Fig. 2, B - E; *t* test, *p* < 0.05). The Simpson index was not altered (Fig. 2, F). The Sobs, Chao, Ace were used to describe the bacterial species richness, *i.e.*, the number of OUTs. The Shannon index and Simpson indexes were used to describe community diversity, including species richness and species evenness. Therefore, deletion of epithelial dicer1 reduces gut bacterial richness and diversity in AD mice.

**Figure 2.**
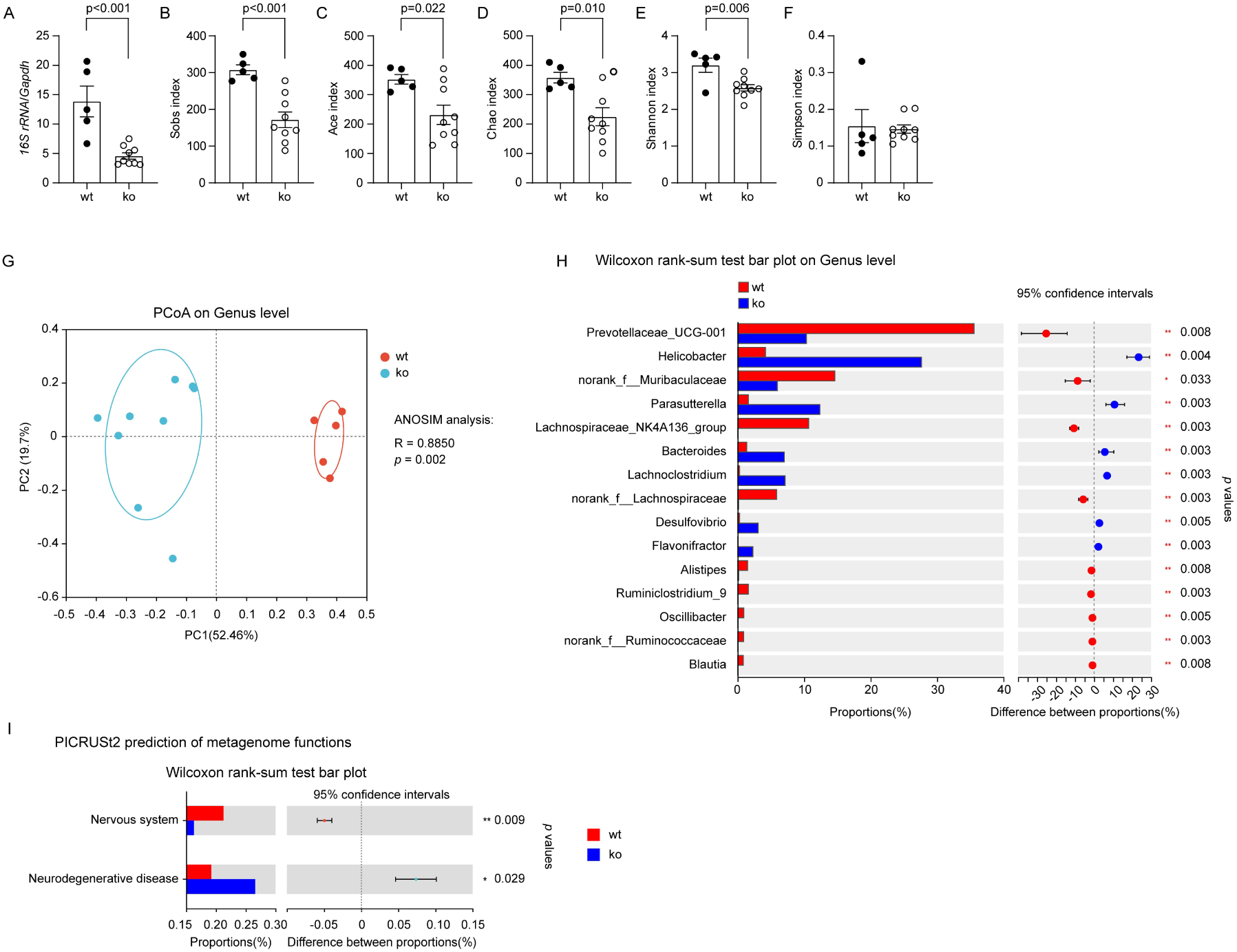
Deletion of dicer1 in gut epithelial cells changed the composition of gut bacteria in *App*-knock-in mice. Nine-month-old *App*-knock-in mice with (ko, n = 9) and without (wt, n = 5) deletion of dicer1 in gut epithelial cells were analyzed for the composition of gut bacteria. Intestinal content for the isolation of bacterial DNA was harvested from the cecum and neighboring colon. Bacterial DNA was first measured for the amount with real-time PCR (A; *t* test), and then sequenced for the V3-V4 region of 16S rRNA-encoding gene. Deletion of dicer1 significantly decreased Sobs, Ace, Chao and Shannon, but not Simpson’s indices (B - F; *t* test), indicating the reduction of bacterial richness and diversity. Principal coordinate analysis (PCoA) was used for β-diversity analysis of bacterial composition at the genus level in *App*-knock-in mice with and without deletion of dicer1 (G; Each symbol represents the gut bacteria of an individual mouse). As expected, the structure of gut bacterial community differed significantly between these two mouse groups (G; ANOSIM analysis). Bar plots depict abundance (% of total) of the indicated genera. Wilcoxon rank-sum tests show that deletion of dicer1 in epithelial cells changed the relative abundance of bacteria in various genera, e.g., *Prevotellaceae*, *Helicobacter* and *Parasutterella* (H). Finally, the PICRUSt2 software was used to predict the function of gut bacteria, which showed that deletion of dicer1 potentially contributes to neurodegenerative disorders (I; Wilcoxon rank-sum tests).

In the PCoA, we clearly observed the difference in the architecture of gut bacteria between App^ki/ki^Dicer1^del^ and App^ki/ki^Dicer1^wt^ mice (Fig. 2, G; ANOSIM, R = 0.8850, *p* = 0.002), which was in line with the observation in mice with constitutive knockout of *Dicer1* gene in intestinal epithelial cells [27]. At the genus level, the abundance of *Helicobacter*, *Parasutterella*, *Bacteroides* and *Lachnoclostrium* increased, while that of *Prevotellaceas*, *Muribaculaceae*, and *Lachnospiraceae* decreased in App^ki/ki^Dicer1^del^ mice compared with App^ki/ki^Dicer1^wt^ controls (Fig. 2, H; Wilcoxon rank-sum test, *p* < 0.01).

To investigate the impact of miRNA deficiency on the brain, we used the PICRUSt2 software [34] and annotated the metagenomic reads on the KEGG database [35] to predict the function of changed microbiota. Deletion of gut miRNA had the potential to influence various pathophysiological functions of the host. Notably, it might promote neurodegenerative disease and adversely affect the nervous system compared with dicer1 wild-type AD mice (Fig. 2, I; Wilcoxon rank-sum test, *p* < 0.05), although the prediction needs to be validated.

In an additional experiment, we pooled all mice and compared the bacterial composition between male and female mice by PCoA analysis. We observed that sex had no effect on the β-diversity of gut bacteria in our App^ki/ki^ mice, which was consistent with our preliminary experiment on the effect of sex on cerebral Aβ levels (see Supplementary Figure 1).

### Deletion of dicer1 in gut epithelial cells modifies microbiome in brain tissues of *App*-knock-in mice

Our recent study has shown that bacterial components, *i.e.*, bacterial DNA, can circulate to the brain [13]. We detected a lower DNA amount of the bacterial 16S rRNA gene in the brain tissue of 9-month-old App^ki/ki^Dicer1^del^ mice compared with App^ki/ki^Dicer1^wt^ littermates (Fig. 3, A; *t* test, *p* < 0.05), which is consistent with bacterial reduction in the gut (Fig. 2, A). Alpha-diversity analysis showed that deletion of dicer1 reduced both the Chao1 index and the observed features (Fig. 3, B and C; *t* test, *p* < 0.05), but did not significantly affect Shannon and Simpson indexes (data not shown). Chao1 index and observed features were positively and strongly correlated (Fig. 3, D; Spearman correlation test, *p* < 0.001). Thus, depletion of miRNA in the gut has the potential to increase the richness and diversity of bacteria in the brain, even if it decreases the absolute number of bacteria.

**Figure 3.**
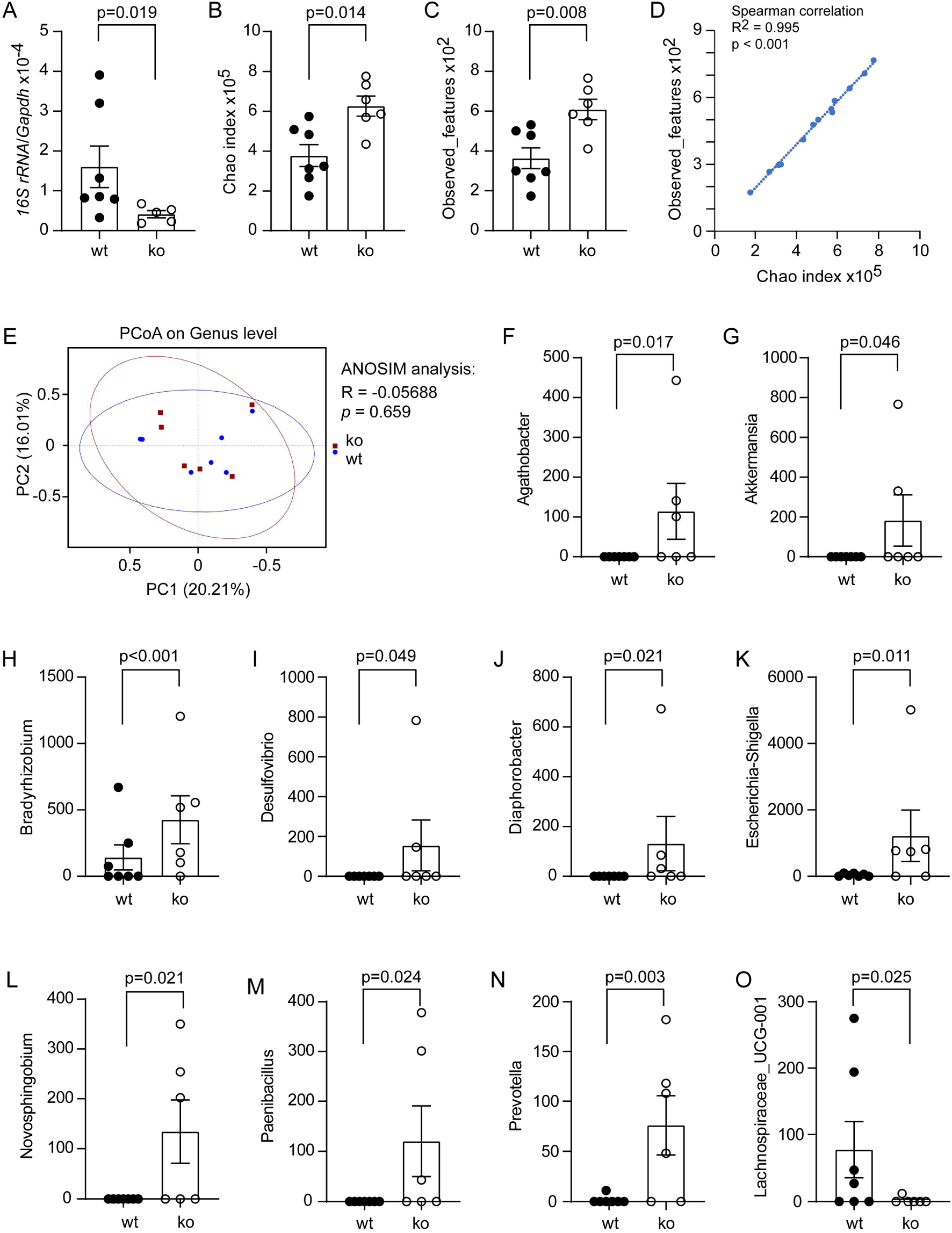
Deletion of dicer1 in gut epithelial cells modified microbiome in the brain of *App*-knock-in mice. Total DNA was isolated from the brain tissues of 9-month-old *App*-knock-in mice with (ko) and without (wt) dicer1 deletion in the gut epithelial cells, measured for the amount of bacterial 16S rDNA and host *Gapdh* DNA with real-time PCR (A; *t* test, *n* = 5 - 7 per group), and then sequenced for the V3-V4 region of 16S rDNA for the microbiome analysis (B - O; *n* = 7 and 6 for wt and ko mice, respectively). Deletion of dicer1 significantly increased Chao index and observed features (B and C; *t* test). Chao index and observed features were strongly correlated (D; Spearman correlation test). Principal coordinate analysis (PCoA) did not show difference in β-diversity analysis of bacterial composition at the genus level in wt and ko *App*-knock-in mice (E; Each symbol represents the cerebral bacteria of an individual mouse; ANOSIM analysis). Bar plots show the abundance of indicated genera in MetagenomeSeq analysis (F - O).

In the brains of 9-month-old App^ki/ki^Dicer1^del^ and App^ki/ki^Dicer1^wt^ littermate mice, we observed no difference in β-diversity analysis at the genus level as shown by PCoA (Fig. 3, E; ANOSIM analysis, *p* > 0.05). However, MetagenomeSeq analysis had shown that deletion of epithelial dicer1 significantly increased the abundance of *Agathobacter*, *Akkermansia*, *Bradyrhizobium*, *Escherichia-Shigella*, *Desulfovibrio*, *Diaphorobacter*, *Novosphingobium*, *Paenibacillus* and *Prevotella* bacteria, and decreased that of *Lachnospiraceae_UCG-001* bacteria (Fig. 3, F - O; non-parametric tests, *p* < 0.05). These bacteria have been found in the gut in association with AD pathology [41–43], or in the blood or cerebrospinal fluid in correlation with inflammatory lesions [44, 45].

It should be noted that the detection and sequencing of bacterial DNA in the brain is still a great challenge. Most reads in our 16S rDNA sequencing were mapped to DNA sequences from bacteria, e.g., *Sphingomonas* and *Acinetobacter* (see Supplementary Figure 2), which are widely found in the environment (water and soil). We need further experiments to exclude the possibility that the environmental bacteria contaminated our bacterial samples from brain tissues.

### Deletion of gut epithelial dicer1 reduces Aβ load in *App*-knock-in mouse brain

Accumulation of Aβ in the brain is the pathological hallmark of AD patients and leads to neurodegeneration in AD [46]. After observing that deletion of epithelial dicer1 altered the microbiome in both the gut and brain of AD mice, we asked whether miRNA deficiency also altered AD pathology in the brain of App^ki/ki^ mice. Four serial brain sections were stained with human Aβ antibodies. Deletion of epithelial dicer1 significantly reduced Aβ coverage in both the hippocampus and cortex of App^ki/ki^Dicer1^del^ mice compared to App^ki/ki^Dicer1^wt^ littermates (Hippocampus: 10.53 % ± 0.44 % vs. 15.29 % ± 0.55 %, n = 11 -12 per group, and Cortex: 9.11 % ± 0.84 % vs. 13.05 % ± 0.36 %, n = 4 -5 per group; Fig. 4, A - C; *t* test, *p* < 0.01).

**Figure 4.**
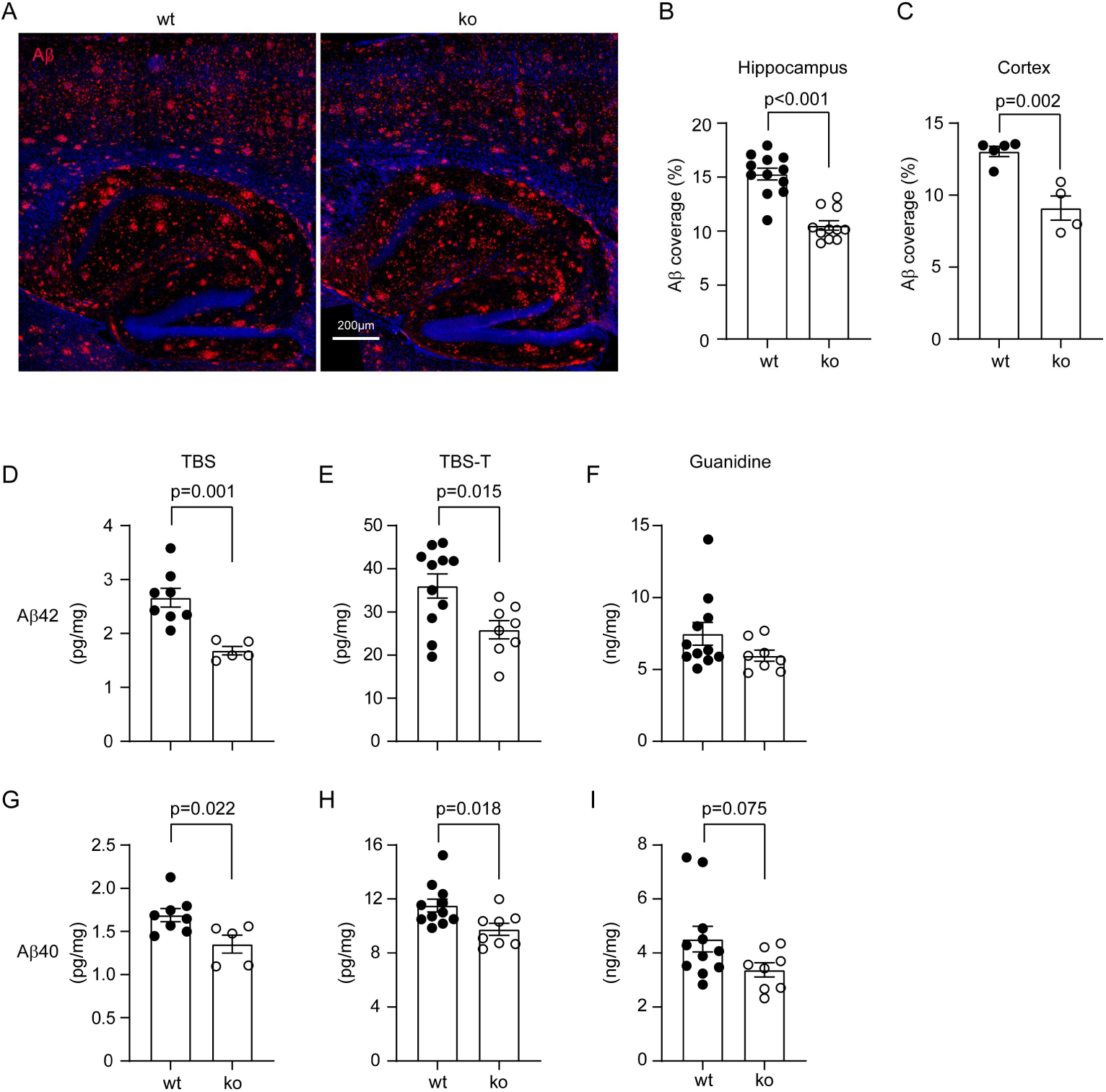
Deletion of dicer1 in gut epithelial cells reduces Aβ load in *App*-knock-in mouse brain. Brains from 9-month-old *App*-knock-in littermate mice with (ko) and without (wt) deletion of epithelial dicer1 were stained for human Aβ (in red) (A). Deletion of dicer1 in epithelial cells significantly decreased Aβ deposits in both the hippocampus and cortex of *App*-knock-in mice (B and C; *t* test; n = 4 - 12 per group). Brain tissues were also serially homogenized and extracted in TBS, TBS plus 1% Triton-100 (TBS-T) and guanidine-HCl, and then measured for Aβ40 and Aβ42 with ELISA. Depletion of gut miRNA decreased the concentrations of both Aβ40 and Aβ42 in TBS- and TBS-T-soluble brain fractions, but not in guanidine-HCl-soluble fraction of AD mice (D - I; *t* test, n = 7 - 9 per group).

We also homogenized brain tissue from 9-month-old tamoxifen-treated App^ki/ki^Dicer1^del^ and App^ki/ki^Dicer1^wt^ mice serially in TBS, TBS-T and guanidine-HCl buffer and measured the concentrations of Aβ42 and Aβ40 in the brain homogenates with ELISA, as we had done previously [37]. We observed that the concentrations of both Aβ42 and Aβ40 in the TBS- and TBS-T-soluble fractions were significantly lower in App^ki/ki^Dicer1^del^ than in App^ki/ki^Dicer1^wt^ mice (Fig. 4, D, E, G and H; *t* test, *p* < 0.05). Deletion of epithelial Dicer1 tended to decrease Aβ40, but not Aβ42 in guanidine-HCl-soluble fractions of brain homogenates (Fig. 4, F and I; *t* test, *p* > 0.05). To exclude the possibility that sex plays a role in the reduction of Aβ40 and Aβ42 induced by miRNA deletion in the gut, we performed a linear regression analysis and confirmed that deletion of epithelial Dicer1 significantly and independently reduced Aβ in the brain (see Table 2; *p* < 0.05). Sex did not affect Aβ levels in the brain, which was consistent with the result of our preliminary experiment (see Supplementary Figure 1).

**Table 2.** Deletion of dicer1 independently modulates cerebral Aβ level.

| Variables |  | Deletion of *dicer1* in gut epithelial cells |  |  | Sex |  |
| --- | --- | --- | --- | --- | --- | --- |
|  |  | Coefficient | *p*-value | 95% confidence interval | Coefficient | *p*-value |
| TBS<br>wt: n = 8 (male: 5)<br>ko: n = 5 (male: 3) | Aβ40 | 0.332 | 0.021 | 0.061 – 0.603 | 0.162 | 0.213 |
|  | Aβ42 | 0.990 | 0.001 | 0.521 – 1.459 | 0.370 | 0.109 |
| TBS-T<br>wt: n = 11 (male: 7)<br>ko: n = 8 (male: 4) | Aβ40 | 1.797 | 0.020 | 0.316 – 3.277 | -0.243 | 0.732 |
|  | Aβ42 | 9.743 | 0.022 | 1.609 – 17.877 | 2.915 | 0.458 |

Code for the analysis: Genotype (deletion of *dicer1* = 0, wt = 1); Sex (male = 1, female = 0)

### Deletion of gut epithelial dicer1 reduces BACE1 activity and increases expression of LRP-1 and ABCB1 at the blood-brain barrier in *App*-knock-in mouse brain

We recently found that depletion of bacteria in the gut decreases the activity of BACE1, a rate-limiting enzyme in Aβ production, and increases the expression of LRP1 and ABCB1, which facilitates the efflux of cerebral Aβ across the blood-brain barrier (BBB) [13]. Therefore, we asked whether these Aβ-reducing mechanisms were also present in App^ki/ki^ mice with deletion of intestinal epithelial dicer1. Interestingly, the activity of BACE1, but not γ-secretase was slightly but significantly lower in brain tissues of 9-month-old App^ki/ki^Dicer1^del^ mice than in App^ki/ki^Dicer1^wt^ mice after tamoxifen treatment (Fig. 5, A and B, two-way ANOVA, *p* < 0.05). Deletion of intestinal epithelial dicer1 also significantly increased protein levels of LRP1 and ABCB1 in cerebral capillaries isolated from App^ki/ki^Dicer1^del^ mice compared with App^ki/ki^Dicer1^wt^ mice (Fig. 5, C - E; *t* test, *p* < 0.05). However, deletion of dicer1 did not alter the expression of PDGFRβ or CD13 in the capillaries of App^ki/ki^ mice (Fig. 5, C, F and G; *t* test, *p* > 0.05). PDGFRβ and CD13 are two protein markers of pericytes that decrease in AD brains [47]. Deletion of intestinal epithelial dicer1 appeared not to increase microglial clearance of Aβ in the brain, as CD68-immunoreactive cells around Aβ deposits did not differ between 9-month-old App^ki/ki^Dicer1^del^ and App^ki/ki^Dicer1^wt^ mice (Fig. 5, H and I; *t* test, *p* > 0.05). The expression of CD68 as a lysosomal protein suggests the activity of microglial phagocytosis [13]. Moreover, depletion of intestinal epithelial dicer1 did not affect the transcription of *Neprilysin* and *Ide* genes (Fig. 5, J and K; *t* test, *p* > 0.05), which encode two important enzymes in the brain digesting Aβ extracellularly [48, 49].

**Figure 5.**
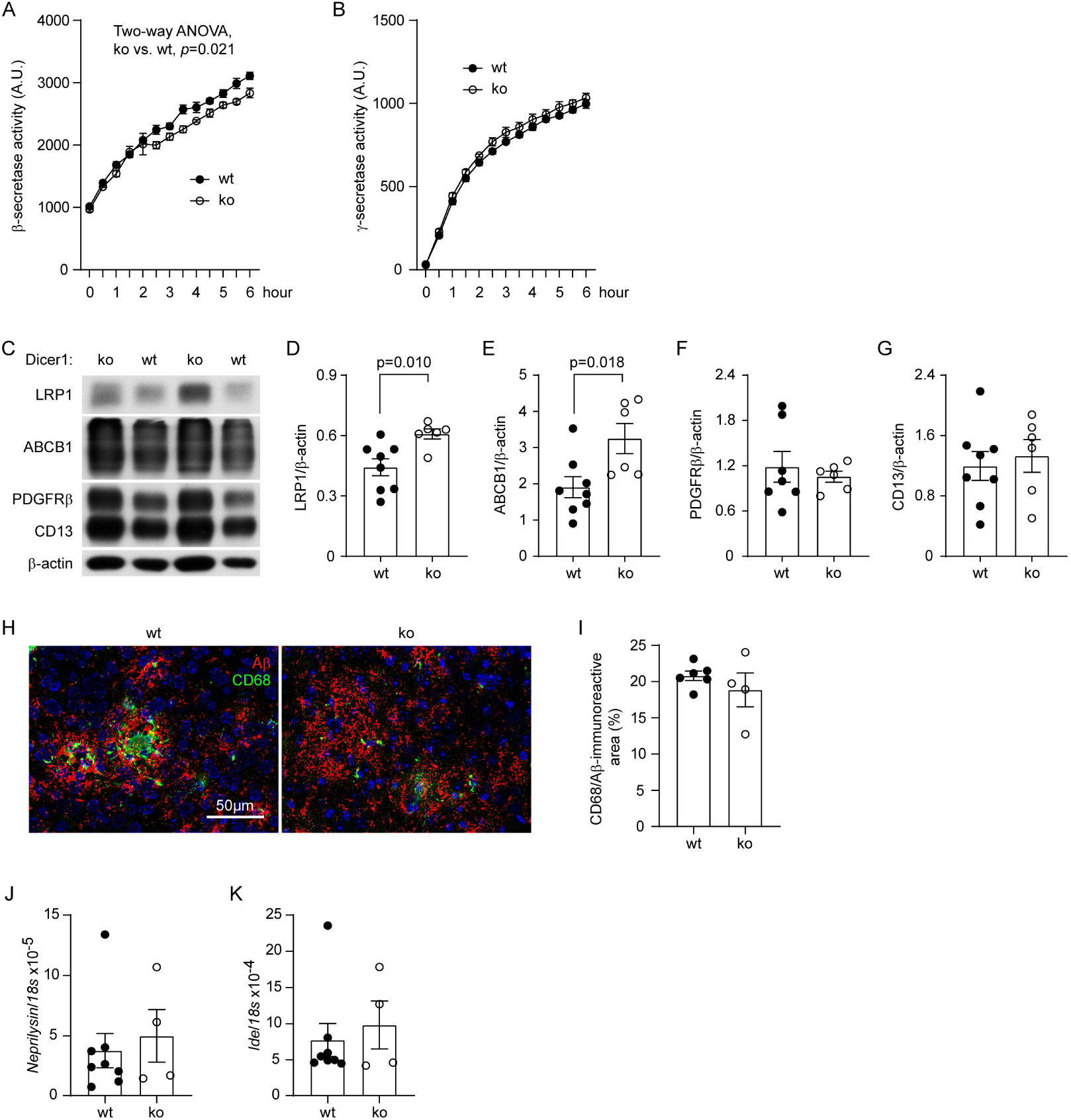
Deletion of dicer1 in gut epithelial cells reduces BACE1 activity and increases LRP1 and ABCB1 expression in the brain of *App*-knock-in mice. Brains from 9-month-old *App*-knock-in littermate mice with (ko) and without (wt) deletion of epithelial dicer1 were prepared for cell membrane components for β- and γ-secretase assays. Deletion of dicer1 slightly but significantly reduced the activity of BACE1, but not γ-secretase (A and B; two-way ANOVA, n = 5 and 3 for wt and ko groups, respectively). Brain tissues were also homogenized for the isolation of capillaries. Quantitative Western blot of vessel lysates showed that deletion of dicer1 significantly increased the protein levels of LRP1 and ABCB1, but not PDGFRβ and CD13 (C - G; *t* test, n = 6 - 8 per group). The brain tissues were further stained with fluorescence-conjugated CD68 and Aβ antibodies (H). Deletion of dicer1 did not change the density of CD68-immunofluorescence around Aβ deposits in both cortex and hippocampus of *App*-knock-in mice (I; *t* test, *n* = 4 - 6 per group). Finally, the transcription of *Neprilysin* and *Ide* genes was determined by real-time PCR, which showed no effect of gut miRNA depletion on the expression of extracellular Aβ-degrading enzymes (J and K; *t* test, *n* = 4 - 8 per group).

### Deletion of gut epithelial dicer1 mildly alter inflammatory activation in the brain of *App*-knock-in mice

Inflammatory activation is a pathological feature of AD in the brain and influences the progression of the disease [50]. In our *App*-knock-in mice, deletion of dicer1 in gut epithelial cells did not alter the numbers of Iba-1-positive microglia in both hippocampus and cortex (Fig. 6, A - C; *t* test, *p* > 0.05). By detecting transcription of individual inflammatory genes, we found that gut miRNA deficiency only significantly reduced *Ccl-2* transcripts and increased the transcription of *Il-10* gene, without altering the transcription of other tested genes (*Tnf-α*, *Il-1β* and *Chi3l3*) (Fig. 6, D - H; *t* test, *p* < 0.05). In the following transcriptomic analysis of brain tissues (Fig. 7), there was no robust difference in inflammatory activation between 9-month-old App^ki/ki^Dicer1^del^ mice and in App^ki/ki^Dicer1^wt^ mice, except that the GO enrichment analysis of differentially expressed genes (DEGs), identified with *p*-value (instead of adjusted *p*-value) cut-off at 0.05 and a minimum absolute log2fold change ≥1, showed that reducing gut miRNA activated the IL-10 signaling pathway in the brain (*p* < 0.05; see Supplementary Figure 3).

**Figure 6.**
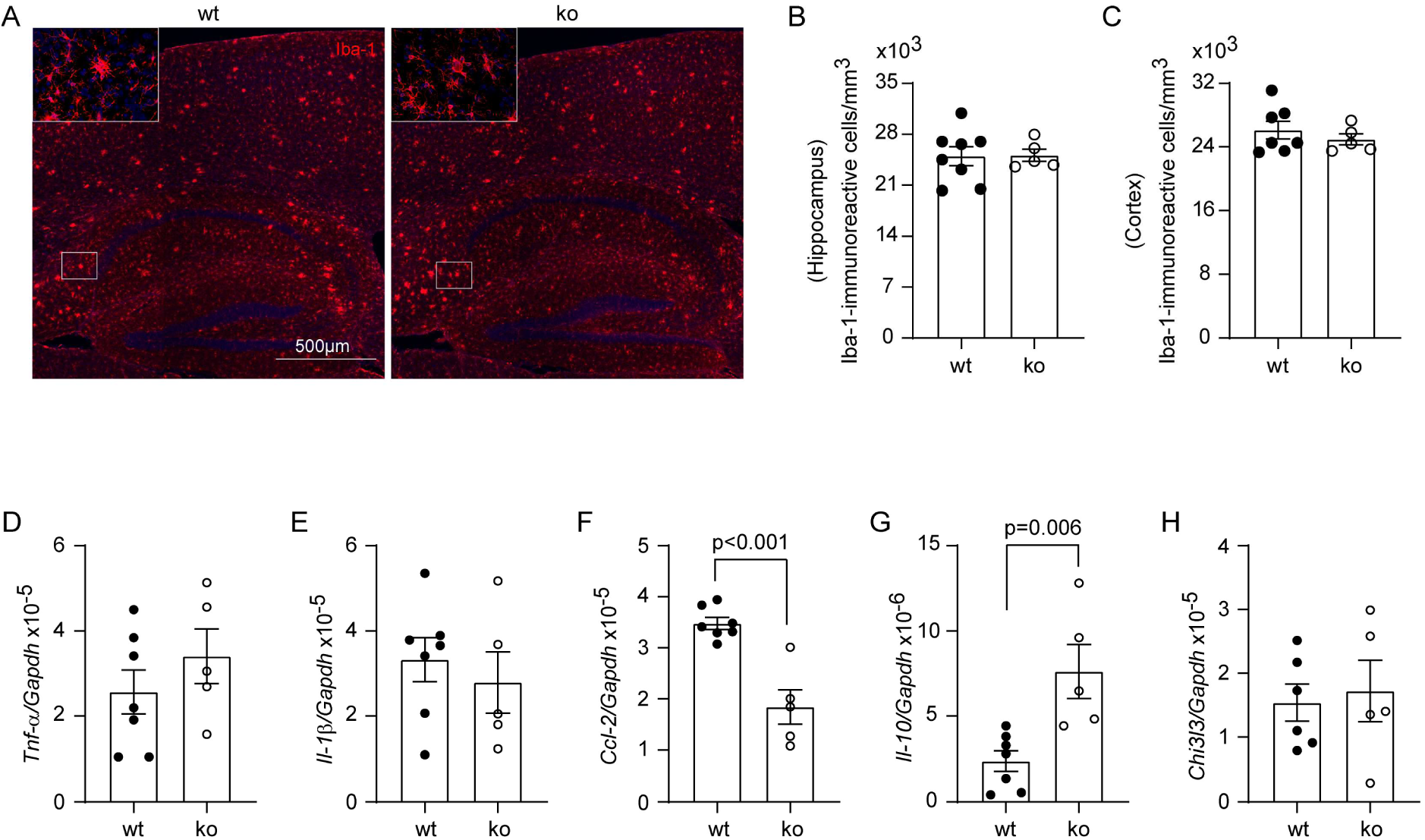
Deletion of dicer1 in gut epithelial cells mildly regulates inflammatory activation in *App*-knock-in mouse brain. Brains from 9-month-old *App*-knock-in littermate mice with (ko) and without (wt) deletion of epithelial dicer1 were stained and counted for Iba-1-positive cells (in red) (A). Deletion of dicer1 in epithelial cells did not change microglial numbers in both the hippocampus and cortex of *App*-knock-in mice (B and C; *t* test; n = 5 - 8 per group). The transcripts of inflammatory genes were also quantified by real-time PCR (D - H). The transcription of *Ccl-2* and *Il-10* genes was down- and upregulated, respectively, in ko compared to wt *App*-knock-in mice (F and G; *t* test, n = 5 - 7 per group).

**Figure 7.**
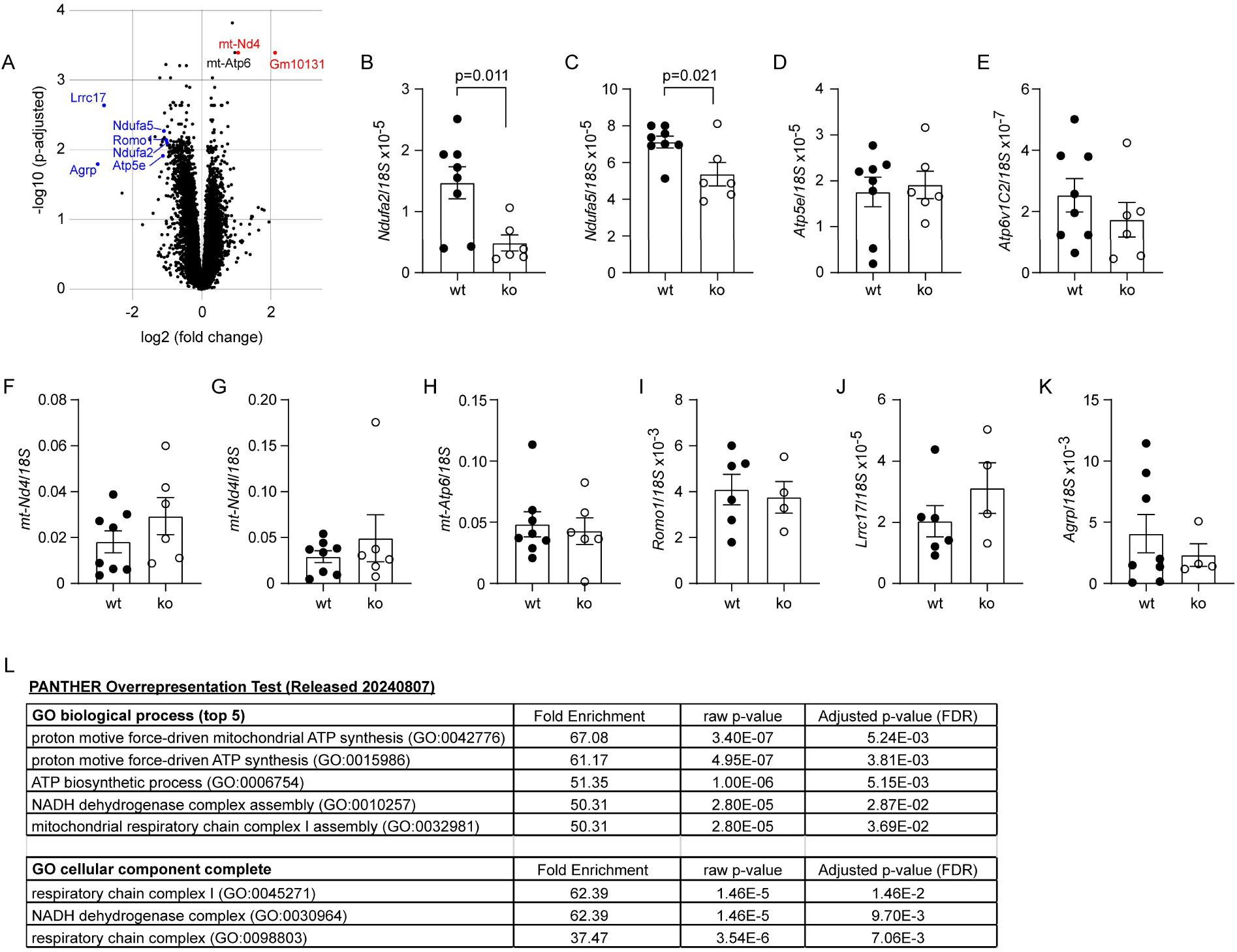
Deletion of dicer1 in gut epithelial cells regulates mitochondrial metabolism-associated gene transcription in *App*-knock-in mouse brain. Total RNA was isolated from the hippocampus and cortex of 9-month-old *App*-knock-in littermate mice with (ko) and without (wt) deletion of epithelial dicer1 for the transcriptomic analysis (*n* = 7 and 12 for ko and wt mice, respectively). Differentially expressed genes (DEGs) were presented in a volcano plot (A) and validated by quantitative RT-PCR (B - K; *t* test; n = 4 - 8 per group). The transcription of *Ndufa2* and *Ndufa5* genes was significantly down-regulated in ko mice compared to wt mice (B and C; *t* test; n = 6 - 8 per group). Thereafter, GO analysis showed that DEGs were associated with mitochondrial energy metabolism (L).

**Figure 8.**
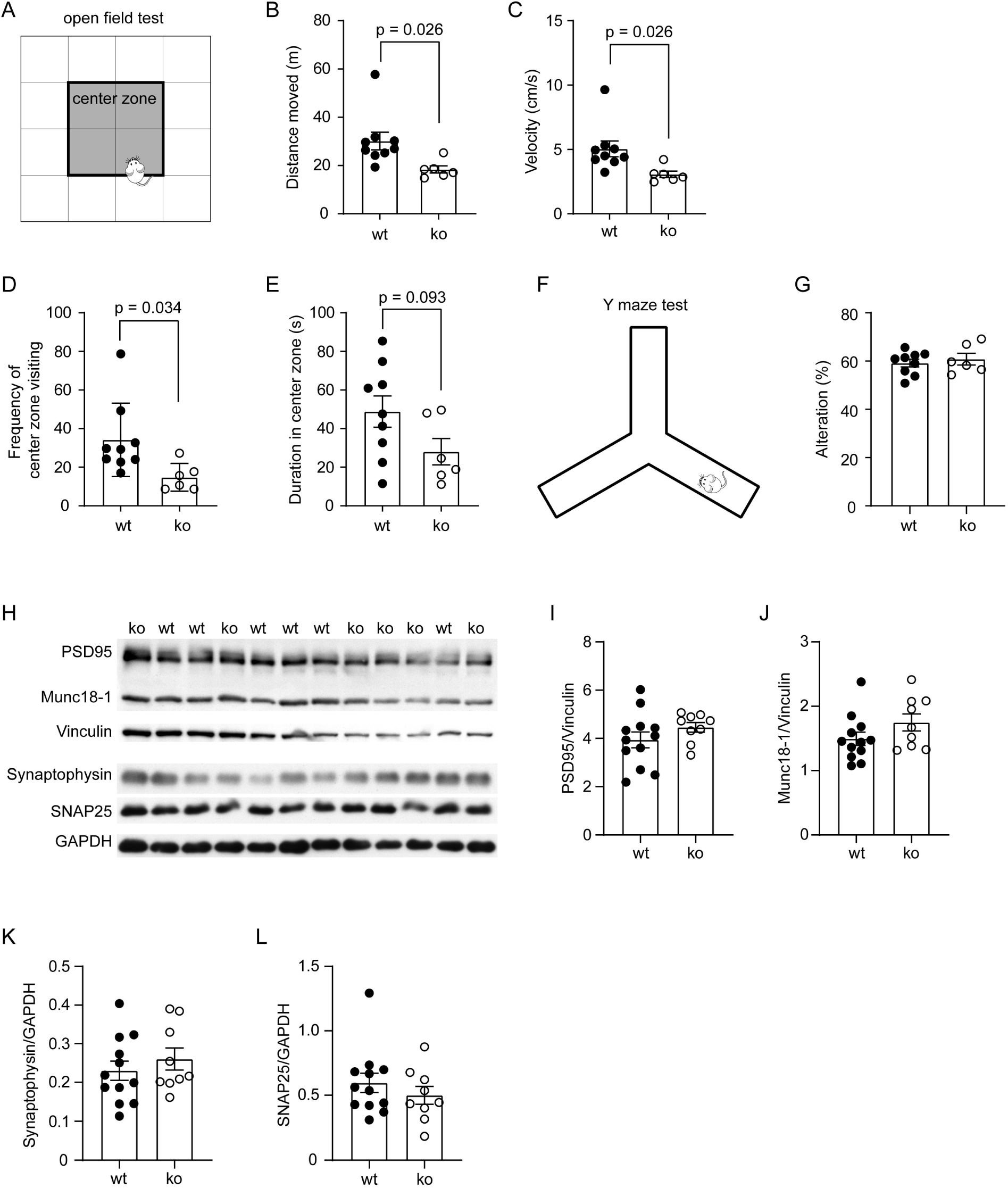
Deletion of dicer1 in gut epithelial cells potentially induces anxiety without improving cognitive function of *App*-knock-in mice. Nine-month-old *App*-knock-in littermate mice with (ko) and without (wt) deletion of epithelial dicer1 received open-field-test (A - E) and Y-maze-test (F and G). Deletion of dicer1 significantly reduced travelling distance and velocity, and the frequency of entries into the center zone of the open field (B - E; *t* test, *n* = 9 and 6 for wt and ko groups, respectively). In Y-maze-test, there was no difference in the alterations between ko and wt mice (G; *t* test, *n* = 9 and 6 for wt and ko groups, respectively). Brain tissues were also homogenized for detection of synaptic proteins by Western blot (H - L). Deletion of dicer1 did not significantly change the protein levels of PSD95, Munc18-1, synaptophysin and SNAP25 in the brain (I - L; *t* test, *n* = 12 and 9 for wt and ko groups, respectively).

### Deletion of gut epithelial dicer1 potentially regulates mitochondrial energy metabolism in the brain of *App*-knock-in mice

To investigate the effects of gut miRNA on the brain systemically, we performed a transcriptome analysis of the hippocampus and cortex of 9-month-old App^ki/ki^Dicer1^del^ and App^ki/ki^Dicer1^wt^ mice. Next-generation sequencing identified 15549 genes, of which 21 and 2 genes were significantly altered by a log2 fold change < -1 and >1 (adjusted *p* < 0.05) in transcription in brain tissue from App^ki/ki^Dicer1^del^ mice compared to App^ki/ki^Dicer1^wt^ mice (Fig. 7, A). Interestingly, most of these DEGs involved mitochondrial metabolism. The only characterized gene with upregulated transcription was a mitochondrial gene *mt-DN4*; and the genes with down-regulated transcription were located in the nucleus and included *Ndufa2* and *Ndufa5*, encoding NADH dehydrogenase [ubiquinone] 1 alpha subunit 2 and 5, respectively, in mitochondrial complex I [51, 52], *Atp5e*, encoding a subunit of ATP synthase, also known as complex V [53], *Romo1*, encoding reactive oxygen species modulator 1 at the inner mitochondrial membrane, which recognizes reactive oxygen species and regulates mitochondrial dynamics [54], and *Agrp*, encoding Agouti-related protein, which increases appetite and decreases metabolism and energy expenditure [55]. Using real-time PCR, we validated that the transcript levels of *Ndufa2* and *Ndufa5*, but not other DEGs, e.g., *Atp5e*, *Atp6v1c2*, *mt-Nd4*, *mt-Nd4l*, *mt-Atp6*, *Romo1*, *Lrrc17*, and *Agrp*, were significantly downregulated in the brains of 9-month-old App^ki/ki^Dicer1^del^ mice compared to App^ki/ki^Dicer1^wt^ mice (Fig. 7, B - K; *t* test, *p* < 0.05). Not surprisingly, GO enrichment analysis of DEGs predicted that depletion of miRNAs in the gut has the potential to reduce ATP production by the mitochondrial respiratory chain (Fig. 7, L; adjusted *t* test, *p* < 0.05).

### Deletion of gut epithelial dicer1 did not improve the cognitive function but might induce anxiety of *App*-knock-in mice

After finding that deletion of intestinal epithelial *Dicer1* gene reduced cerebral Aβ and altered neuroinflammation in App^ki/ki^ mice, we further analyzed the effects of intestinal miRNA on neuronal function. We performed open field test to probe anxiety and Y-maze test for the cognitive function in 9-month-old tamoxifen-treated App^ki/ki^Dicer1^del^ and App^ki/ki^Dicer1^wt^ mice. App^ki/ki^ mice have been reported to display anxiety-related symptoms, even before the occurrence of cognitive dysfunction [56, 57]. In the open field test, App^ki/ki^Dicer1^del^ mice travelled significantly shorter distance and at a lower speed than App^ki/ki^Dicer1^wt^ mice (Fig. 8, A - C; *t* test, *p* < 0.05); while, we did not observe any problems in motor function of App^ki/ki^Dicer1^del^ mice (data not shown). We subsequently divided the arena into an outer zone and a center zone (Fig. 8, A) and observed that deletion of epithelial *Dicer1* gene significantly reduced the entries to the center zone (Fig. 8, D; *t* test, *p* < 0.05) and tendered to decrease the traveling duration in the center area (Fig. 8, E; *t* test, *p* = 0.093), suggesting an anxiogenic phenotype. In Y-maze test, App^ki/ki^Dicer1^del^ and App^ki/ki^Dicer1^wt^ mice did not differ in the alteration of arm entries (Fig. 8, F and G; *t* test, *p* > 0.05), indicating that deletion of intestinal miRNA did not improve the cognitive function.

We also homogenized brains from 9-month-old App^ki/ki^Dicer1^del^ and App^ki/ki^Dicer1^wt^ mice and determined synaptic proteins with quantitative Western blot (Fig. 8, H). Deletion of epithelial *Dicer1* gene did not alter the protein levels of PSD-95, Munc18-1, synaptophysin and SNAP25 (Fig. 8, I - L; *t* test, *p* > 0.05), which was in accordance with the results of the Y-maze test.

## Discussion

There is increasing evidence that the gut microbiome is altered in AD patients and animal models, which is associated with AD pathologies in the brain [3–7, 11–13, 17]. However, the gut microbiome is very different between AD patients or between AD animals, so AD-specific profile of the gut microbiome cannot be defined. The potential therapy targeting gut bacteria may need to be personalized. The aim of our study was to develop a miRNA-based therapeutic method for the flexible, efficient and specific modification of AD-associated bacterial taxon in the gut. The current exploratory project has shown that knockout of the gene *Dicer1*, encoding miRNA-producing enzyme, in intestinal epithelial cells alters both the gut and brain microbiome without compromising the intestinal epithelial barrier, and reduces Aβ pathology in the brain. However, depletion of overall miRNA in the gut has the potential to inhibit mitochondrial energy production in the brain and increase anxiety of *App*-knock-in mice.

In our AD mice, we used inducible Cre recombinase driven by *Villin* gene promoter to delete dicer1 in intestinal epithelial cells. We confirmed the original description of Villin-Cre mice that Cre-mediated gene recombination can persist over many cycles of epithelial renewal and even two months after tamoxifen induction [32]. Due to technical difficulties, we were unable to directly validate the depletion of miRNA in the intestinal lumen. However, changes in gut microbiome and expression of TJPs in the cecum, which were consistent with the previous study on constitutive knockout of *Dicer1* in the intestinal epithelial cells [27], strongly suggested successful depletion of miRNA in the gut of our AD mice. Of note, inducible knockout of *Dicer1* gene in our AD model reduced the transcription of inflammatory genes (e.g. *Tnf-α*, *Il-1β* and *Inos*) in the cecum and did not alter the transcription of *Il-17a*, *Ifn-γ*, *Il-4* and *Il-10* genes in CD4-positive spleen cells, indicating that depletion of miRNA in the intestine is unlikely to impair intestinal barrier function in AD mice. Our results differed from the observation in mice with dextran sodium sulfate [DSS]-induced colitis, in which knockout of *Dicer1* gene increased susceptibility to colitis [27]. The toxic pressure on the intestinal barrier in inflammatory bowel disease is obviously higher than in AD. On the other hand, the inhibition of inflammation in the cecum, which correlates with the reduction in intestinal bacteria, could be another indication of bacteria-mediated effects of miRNA in the gut. Moreover, the inhibition of TJP expression could even be the result of bacterial reduction, as gut bacteria stimulate TJP expression [58]. Therefore, the phenotype of our *Dicer1* knockout AD model is mainly due to the absence of gut miRNA.

Intestinal miRNA has been reported to enter bacteria, interfere with mRNA transcription and change bacterial proliferation and metabolism [27]. Oral treatment with synthetically produced miRNA mimics increases the abundance of *Akkermansia muciniphila* bacteria in the gut, expands Treg lymphocytes and attenuates disease severity in EAE mice [29]. We observed that overall depletion of miRNA in the gut decreased the abundance of bacteria from *Prevotellaceae* UCG-001 and Norank_f_*Muribaculaceae* genera, and increased the proportions of *Helicobacter*, *Parasutterella*, *Bacteroides*, and *Lachnoclostridium* bacteria. Bacteria of the genus *Prevotella* and the family *Muribaculaceae* (formerly known as S24-7) produce short-chain free fatty acids (SCFAs; e.g., butyrate), serving as potential probiotic bacteria [59, 60]. However, direct evidence for the anti-inflammatory and AD-preventing effects of *Prevotellaceae* UCG-001 and Norank_f_*Muribaculaceae* is still pending. Infection of *Helicobacter pylori* has been associated with the risk of AD [61]. *Bacteroides* bacteria frequently increase in the gut of AD patients [62]. However, the pathogenic effects of the changes in *Helicobacter* and *Bacteroides* might be compromised as the absolute number of bacteria in the gut of our miRNA-deficient *App*-knock-in mice decreased.

It is noteworthy that various manipulations of gut bacteria leading to a reduction in α-diversity appear to attenuate cerebral Aβ pathology in AD mice [10, 15, 16, 19, 63]. The same applies to our gut miRNA-depleted *App*-knock-in mice. The α-diversity in the gut microbiome of AD patients often decreases [3, 6, 20]. Whether this is a protective mechanism for slowing the progression of AD remains to be clarified. Our recent studies have shown that changes in gut bacteria affect at least two mechanisms that alter cerebral Aβ levels: 1) inhibition of β-secretase activity, which reduces Aβ production, and 2) increase in the expression of LRP1 and ABCB1 at the blood-brain barrier, both of which mediate the efflux of Aβ from the brain parenchyma into the bloodstream [13]. We have shown that knockout of *Il-17a* gene or depletion of gut bacteria triggers these two mechanisms and attenuates Aβ pathology in APP/PS1-transgenic mice [13, 64]. However, no changes in the transcription of *Il-17a*, as well as *Ifn-γ*, *Il-4* and *Il-10* genes were detected in CD4-positive splenocytes of intestinal miRNA-depleted AD mice, suggesting that there are other molecular signaling pathways than IL-17a inhibition mediating the Aβ-reducing effects. SCFAs potentially increase the activity of β-secretase [10], and act on the blood-brain barrier by promoting expression of TJPs and reducing permeability [65]. We observed in the gut a reduction in both absolute number of bacteria and proportions of *Prevotellaceae* UCG-001, *Norank_f_Muribaculaceae*, *Lachnospiraceae* and *Blautia* bacteria that produce SCFAs. It is plausible that depletion of gut miRNA leads to fewer SCFAs in the brain, although SCFAs were not examined in our mice.

Interestingly, we observed that depletion of miRNA in the gut reduced the cerebral transcription of *Ndufa2* and *Ndufa5* genes, encoding subunits of NADH:ubiquinone oxidoreductase, the first and largest complex of the mitochondrial respiratory chain, which mediates the ATP generation and redox processes. A loss-of-function mutation of *NDUFA2* leads to Leigh disease, which is characterized by neurodegeneration [51]. Knockout of *Ndufa5* gene in neurons impairs the assembly and activity of complex I, leading to mild chronic encephalopathy, although it does not cause oxidative damage, neuronal death or gliosis in the brains of mice [52]. Notably, there was also a study showing that mild inhibition of mitochondrial complex I activates AMPK and inhibits GSK3β activity, which could reduce Aβ and tau phosphorylation, restore axonal transport and prevent cognitive deficits in 5× FAD mice [66]. Therefore, the intricate effects of miRNA and gut microbiota on mitochondria pathology in AD brain need to be further and comprehensively investigated.

AD is classically associated with cognitive decline, but other behavioral changes, such as reduced social engagement and increased anxiety, also occur early in AD patients, even before cognitive dysfunction. Anxiety is common in AD, with up to 71% of patients reporting anxiety concerns [67, 68]. Anxiety might also promote or be related to the conversion from mild cognitive impairment (MCI) to AD. As reported, 83.3% of MCI patients that exhibit anxiety symptoms, whereas only 40.9% of persons with MCI but without anxiety, convert to AD within a 3-year follow-up period [69]. The gut miRNA-depleted *App*-knock-in mice studied in this project showed anxious symptoms in the open field test, indicating that a sufficient amount and diversity of miRNA in the gut is necessary for brain health.

Our study addressed the highly debatable question of whether bacteria can spread in the brain. We detected bacterial DNA in brain tissue, as we did in a recent study [13]. In both studies, we found that the reduction of bacterial DNA in the brain correlated with the reduction of bacteria in the gut, suggesting that the bacterial DNA in the brain may partially originate from the gut. However, we still need to confirm the same identity of the bacterial DNA in the brain and gut, and investigate how the bacterial DNA circulates from the gut to the brain. It is still a big challenge to detect and characterize bacteria in the brain. The mitochondrial DNA from the host interferes with the sequencing of bacterial DNA. It is difficult to avoid contamination of experimental samples by environmental bacteria. In previous studies, bacterial DNA detected in the brain, which is identical to bacterial DNA from the environment, was often regarded as sample contamination [70–72]. In fact, this conclusion would need to be validated by further experiments. In our current study, all our tissue samples isolated from AD mice were performed by the same experimenter in the same laboratory and at the same time, so the probability of contamination was the same. Our observation of the difference in bacterial DNA in the brains of dicer1-deficient and wild-type *App*-knock-in mice should not be a mere artifact. We observed that deletion of dicer1 in the intestinal epithelial cells increased the abundance in the brain of some typical gut bacteria, e.g., *Akkermansia*, *Prevotella* and *Escherichia-Shigella*. These bacteria in the brain should have different effects than those in the gut, as the environment and nutrient supply for the bacteria in the brain have changed. *Prevotella*, as *Porphyromonas gingivalis*, is an important pathogen of chronic periodontitis [73]. *P. gingivalis* has been identified in the AD brain and contributes to AD pathogenesis [74]. However, it should be reiterated that the effects of these bacteria in the brain could be attenuated by reducing their absolute numbers in dicer1-deficient *App*-knock-in mice. The most important question for the future will therefore be what biological significance the presence of bacteria (regardless of living cells or cellular components) in the brain has for AD development.

## Conclusions

In summary, depletion of miRNA by conditional knockout of *Dicer1* gene in intestinal epithelial cells reduces α-diversity and alters the composition of bacteria in the gut, and perhaps also in the brain. Depletion of miRNA in the gut attenuates Aβ pathology but may also inhibit mitochondrial respiration in the brain of AD mice. Reduction of miRNA could induce anxiety of AD mice as well. An obvious limitation of our study is that we non-specifically depleted miRNA in the gut. Nevertheless, our study has demonstrated a novel pathway that oral administration of miRNA mimics has the potential to modulate AD pathogenesis in the brain. Future studies should focus on the identification of AD-specific miRNAs in the gut that can be therapeutically utilized to alter the gut microbiome and prevent AD progression.

## Data availability statement

The datasets of the transcriptome and microbiome in brain tissue generated in our study are deposited in Figshare (https://doi.org/10.6084/m9.figshare.28904348.v1).

## Acknowledgments

We thank Dr. A. Domanskyi (University of Helsinki) for providing *Dicer1*-floxed mice, Dr. E. Batlle (The Barcelona Institute of Science and Technology) for Villin-CreERT2 transgenic mice, and Dr. T.C. Saido (RIKEN Brain Science Institute) for *App^NL-G-F^* knock-in mice through the RIKEN Bioresource Center, RIKEN Tsukuba Institute, Japan. We appreciate Elisabeth Gluding and Kati Jordan for their excellent technical assistance.

## Author contributions

Y.L. conceptualized and designed the study, acquired funding, conducted experiments, acquired and analyzed data, and wrote the manuscript. W.H., Q.L., I.S., G.G., S.T., W.Q., H.C., and G.W. conducted experiments, acquired data and analyzed data. S.G. and J.S. provided research laboratories and supervised the research work. All authors contributed to the article and approved the submitted version.

## Ethics declarations

Ethics approval and consent to participate

Animal experiments were conducted in accordance with national rules and ARRIVE guidelines, and authorized by Landesamt für Verbraucherschutz, Saarland, Germany (registration numbers: 36/2018).

## Consent for publication

Not applicable.

## References

1. Fung TC, Olson CA, Hsiao EY: Interactions between the microbiota, immune and nervous systems in health and disease. Nat Neurosci 2017, 20(2):145–155.

2. Gershon MD, Margolis KG: The gut, its microbiome, and the brain: connections and communications. J Clin Invest 2021, 131(18):e143768.

3. Murray ER, Kemp M, Nguyen TT: The Microbiota-Gut-Brain Axis in Alzheimer’s Disease: A Review of Taxonomic Alterations and Potential Avenues for Interventions. Arch Clin Neuropsychol 2022, 37(3):595–607.

4. Ferreiro AL, Choi J, Ryou J, Newcomer EP, Thompson R, Bollinger RM, Hall-Moore C, Ndao IM, Sax L, Benzinger TLS et al: Gut microbiome composition may be an indicator of preclinical Alzheimer’s disease. Sci Transl Med 2023, 15(700):eabo2984.

5. Cattaneo A, Cattane N, Galluzzi S, Provasi S, Lopizzo N, Festari C, Ferrari C, Guerra UP, Paghera B, Muscio C et al: Association of brain amyloidosis with pro-inflammatory gut bacterial taxa and peripheral inflammation markers in cognitively impaired elderly. Neurobiol Aging 2017, 49:60–68.

6. Vogt NM, Kerby RL, Dill-McFarland KA, Harding SJ, Merluzzi AP, Johnson SC, Carlsson CM, Asthana S, Zetterberg H, Blennow K et al: Gut microbiome alterations in Alzheimer’s disease. Sci Rep 2017, 7(1):13537.

7. Verhaar BJH, Hendriksen HMA, de Leeuw FA, Doorduijn AS, van Leeuwenstijn M, Teunissen CE, Barkhof F, Scheltens P, Kraaij R, van Duijn CM et al: Gut Microbiota Composition Is Related to AD Pathology. Front Immunol 2021, 12:794519.

8. Rakusa E, Fink A, Tamguney G, Heneka MT, Doblhammer G: Sporadic Use of Antibiotics in Older Adults and the Risk of Dementia: A Nested Case-Control Study Based on German Health Claims Data. J Alzheimers Dis 2023, 93(4):1329–1339.

9. Grabrucker S, Marizzoni M, Silajdzic E, Lopizzo N, Mombelli E, Nicolas S, Dohm-Hansen S, Scassellati C, Moretti DV, Rosa M et al: Microbiota from Alzheimer’s patients induce deficits in cognition and hippocampal neurogenesis. Brain 2023, 146(12):4916–4934.

10. Colombo AV, Sadler RK, Llovera G, Singh V, Roth S, Heindl S, Sebastian Monasor L, Verhoeven A, Peters F, Parhizkar S et al: Microbiota-derived short chain fatty acids modulate microglia and promote Abeta plaque deposition. Elife 2021, 10:e59826.

11. Xie J, Bruggeman A, De Nolf C, Vandendriessche C, Van Imschoot G, Van Wonterghem E, Vereecke L, Vandenbroucke RE: Gut microbiota regulates blood-cerebrospinal fluid barrier function and Abeta pathology. EMBO J 2023, 42(17):e111515.

12. Erny D, Dokalis N, Mezo C, Castoldi A, Mossad O, Staszewski O, Frosch M, Villa M, Fuchs V, Mayer A et al: Microbiota-derived acetate enables the metabolic fitness of the brain innate immune system during health and disease. Cell Metab 2021, 33(11):2260–2276 e2267.

13. Hao W, Luo Q, Tomic I, Quan W, Hartmann T, Menger MD, Fassbender K, Liu Y: Modulation of Alzheimer’s disease brain pathology in mice by gut bacterial depletion: the role of IL-17a. Gut Microbes 2024, 16(1):2363014.

14. Huang Y, Wu J, Zhang H, Li Y, Wen L, Tan X, Cheng K, Liu Y, Pu J, Liu L et al: The gut microbiome modulates the transformation of microglial subtypes. Mol Psychiatry 2023, 28(4):1611–1621.

15. Minter MR, Hinterleitner R, Meisel M, Zhang C, Leone V, Zhang X, Oyler-Castrillo P, Zhang X, Musch MW, Shen X et al: Antibiotic-induced perturbations in microbial diversity during post-natal development alters amyloid pathology in an aged APPSWE/PS1DeltaE9 murine model of Alzheimer’s disease. Sci Rep 2017, 7(1):10411.

16. Minter MR, Zhang C, Leone V, Ringus DL, Zhang X, Oyler-Castrillo P, Musch MW, Liao F, Ward JF, Holtzman DM et al: Antibiotic-induced perturbations in gut microbial diversity influences neuro-inflammation and amyloidosis in a murine model of Alzheimer’s disease. Sci Rep 2016, 6:30028.

17. Harach T, Marungruang N, Duthilleul N, Cheatham V, Mc Coy KD, Frisoni G, Neher JJ, Fak F, Jucker M, Lasser T et al: Reduction of Abeta amyloid pathology in APPPS1 transgenic mice in the absence of gut microbiota. Sci Rep 2017, 7:41802.

18. Kaur H, Nookala S, Singh S, Mukundan S, Nagamoto-Combs K, Combs CK: Sex-Dependent Effects of Intestinal Microbiome Manipulation in a Mouse Model of Alzheimer’s Disease. Cells 2021, 10(9):2370.

19. Dodiya HB, Kuntz T, Shaik SM, Baufeld C, Leibowitz J, Zhang X, Gottel N, Zhang X, Butovsky O, Gilbert JA et al: Sex-specific effects of microbiome perturbations on cerebral Abeta amyloidosis and microglia phenotypes. J Exp Med 2019, 216(7):1542–1560.

20. Liu P, Wu L, Peng G, Han Y, Tang R, Ge J, Zhang L, Jia L, Yue S, Zhou K et al: Altered microbiomes distinguish Alzheimer’s disease from amnestic mild cognitive impairment and health in a Chinese cohort. Brain Behav Immun 2019, 80:633–643.

21. Li B, He Y, Ma J, Huang P, Du J, Cao L, Wang Y, Xiao Q, Tang H, Chen S: Mild cognitive impairment has similar alterations as Alzheimer’s disease in gut microbiota. Alzheimers Dement 2019, 15(10):1357–1366.

22. Haran JP, Bhattarai SK, Foley SE, Dutta P, Ward DV, Bucci V, McCormick BA: Alzheimer’s Disease Microbiome Is Associated with Dysregulation of the Anti-Inflammatory P-Glycoprotein Pathway. mBio 2019, 10(3):e00632–00619.

23. Vital M, Karch A, Pieper DH: Colonic Butyrate-Producing Communities in Humans: an Overview Using Omics Data. mSystems 2017, 2(6):e00130–00117.

24. Singh V, Lee G, Son H, Koh H, Kim ES, Unno T, Shin JH: Butyrate producers, “The Sentinel of Gut”: Their intestinal significance with and beyond butyrate, and prospective use as microbial therapeutics. Front Microbiol 2022, 13:1103836.

25. Zhang T, Gao G, Kwok LY, Sun Z: Gut microbiome-targeted therapies for Alzheimer’s disease. Gut Microbes 2023, 15(2):2271613.

26. Chilton PM, Ghare SS, Charpentier BT, Myers SA, Rao AV, Petrosino JF, Hoffman KL, Greenwell JC, Tyagi N, Behera J et al: Age-associated temporal decline in butyrate-producing bacteria plays a key pathogenic role in the onset and progression of neuropathology and memory deficits in 3×Tg-AD mice. Gut Microbes 2024, 16(1):2389319.

27. Liu S, da Cunha AP, Rezende RM, Cialic R, Wei Z, Bry L, Comstock LE, Gandhi R, Weiner HL: The Host Shapes the Gut Microbiota via Fecal MicroRNA. Cell Host Microbe 2016, 19(1):32–43.

28. Lv Y, Zhen C, Liu A, Hu Y, Yang G, Xu C, Lou Y, Cheng Q, Luo Y, Yu J et al: Profiles and interactions of gut microbiome and intestinal microRNAs in pediatric Crohn’s disease. mSystems 2024:e0078324.

29. Liu S, Rezende RM, Moreira TG, Tankou SK, Cox LM, Wu M, Song A, Dhang FH, Wei Z, Costamagna G et al: Oral Administration of miR-30d from Feces of MS Patients Suppresses MS-like Symptoms in Mice by Expanding Akkermansia muciniphila. Cell Host Microbe 2019, 26(6):779–794 e778.

30. Saito T, Matsuba Y, Mihira N, Takano J, Nilsson P, Itohara S, Iwata N, Saido TC: Single App knock-in mouse models of Alzheimer’s disease. Nat Neurosci 2014, 17(5):661–663.

31. Cobb BS, Nesterova TB, Thompson E, Hertweck A, O’Connor E, Godwin J, Wilson CB, Brockdorff N, Fisher AG, Smale ST et al: T cell lineage choice and differentiation in the absence of the RNase III enzyme Dicer. J Exp Med 2005, 201(9):1367–1373.

32. el Marjou F, Janssen KP, Chang BH, Li M, Hindie V, Chan L, Louvard D, Chambon P, Metzger D, Robine S: Tissue-specific and inducible Cre-mediated recombination in the gut epithelium. Genesis 2004, 39(3):186–193.

33. Quan W, Luo Q, Hao W, Tomic I, Furihata T, Schulz-Schaffer W, Menger MD, Fassbender K, Liu Y: Haploinsufficiency of microglial MyD88 ameliorates Alzheimer’s pathology and vascular disorders in APP/PS1-transgenic mice. Glia 2021, 69(8):1987–2005.

34. Douglas GM, Maffei VJ, Zaneveld JR, Yurgel SN, Brown JR, Taylor CM, Huttenhower C, Langille MGI: PICRUSt2 for prediction of metagenome functions. Nature Biotechnology 2020, 38(6):685–688.

35. Kanehisa M, Goto S: KEGG: kyoto encyclopedia of genes and genomes. Nucleic Acids Res 2000, 28(1):27–30.

36. Paulson JN, Stine OC, Bravo HC, Pop M: Differential abundance analysis for microbial marker-gene surveys. Nature Methods 2013, 10(12):1200–1202.

37. Liu Y, Liu X, Hao W, Decker Y, Schomburg R, Fulop L, Pasparakis M, Menger MD, Fassbender K: IKKbeta deficiency in myeloid cells ameliorates Alzheimer’s disease-related symptoms and pathology. J Neurosci 2014, 34(39):12982–12999.

38. Hao W, Decker Y, Schnoder L, Schottek A, Li D, Menger MD, Fassbender K, Liu Y: Deficiency of IkappaB Kinase beta in Myeloid Cells Reduces Severity of Experimental Autoimmune Encephalomyelitis. Am J Pathol 2016, 186(5):1245–1257.

39. Liu W, Chen Y, Golan MA, Annunziata ML, Du J, Dougherty U, Kong J, Musch M, Huang Y, Pekow J et al: Intestinal epithelial vitamin D receptor signaling inhibits experimental colitis. J Clin Invest 2013, 123(9):3983–3996.

40. El-Merhie N, Baumgart-Vogt E, Pilatz A, Pfreimer S, Pfeiffer B, Pak O, Kosanovic D, Seimetz M, Schermuly RT, Weissmann N et al: Differential Alterations of the Mitochondrial Morphology and Respiratory Chain Complexes during Postnatal Development of the Mouse Lung. Oxid Med Cell Longev 2017, 2017:9169146.

41. Liu GS, Song Y, Yan JS, Chai YJ, Zhao YF, Ma H: Identification of enterotype for patients with Alzheimer’s disease. J Transl Med 2025, 23(1):299.

42. Lwere K, Muwonge H, Sendagire H, Sajatovic M, Williams SM, Gumukiriza-Onoria JL, Buwembo D, Buwembo W, Nassanga R, Nakimbugwe R et al: Characterization of the gut microbiome in Alzheimer disease and mild cognitive impairment among older adults in Uganda: A case-control study. Medicine (Baltimore) 2025, 104(16):e42100.

43. Chevalier C, Tournier BB, Marizzoni M, Park R, Paquis A, Ceyzeriat K, Badina AM, Lathuiliere A, Saleri S, Cillis F et al: Fecal Microbiota Transplantation (FMT) From a Human at Low Risk for Alzheimer’s Disease Improves Short-Term Recognition Memory and Increases Neuroinflammation in a 3xTg AD Mouse Model. Genes Brain Behav 2025, 24(1):e70012.

44. Gedgaudas R, Bajaj JS, Skieceviciene J, Varkalaite G, Jurkeviciute G, Gelman S, Valantiene I, Zykus R, Pranculis A, Bang C et al: Circulating microbiome in patients with portal hypertension. Gut Microbes 2022, 14(1):2029674.

45. Morton SU, Hehnly C, Burgoine K, Ssentongo P, Ericson JE, Kumar MS, Hagmann C, Fronterre C, Smith J, Movassagh M et al: *Paenibacillus* spp infection among infants with postinfectious hydrocephalus in Uganda: an observational case-control study. The Lancet Microbe 2023, 4(8):e601–e611.

46. Scheltens P, De Strooper B, Kivipelto M, Holstege H, Chetelat G, Teunissen CE, Cummings J, van der Flier WM: Alzheimer’s disease. Lancet 2021, 397(10284):1577–1590.

47. Ding R, Hase Y, Ameen-Ali KE, Ndung’u M, Stevenson W, Barsby J, Gourlay R, Akinyemi T, Akinyemi R, Uemura MT et al: Loss of capillary pericytes and the blood-brain barrier in white matter in poststroke and vascular dementias and Alzheimer’s disease. Brain Pathol 2020, 30(6):1087–1101.

48. Leissring MA, Farris W, Chang AY, Walsh DM, Wu X, Sun X, Frosch MP, Selkoe DJ: Enhanced proteolysis of beta-amyloid in APP transgenic mice prevents plaque formation, secondary pathology, and premature death. Neuron 2003, 40(6):1087–1093.

49. Miners JS, Baig S, Palmer J, Palmer LE, Kehoe PG, Love S: Abeta-degrading enzymes in Alzheimer’s disease. Brain Pathol 2008, 18(2):240–252.

50. Heneka MT, van der Flier WM, Jessen F, Hoozemanns J, Thal DR, Boche D, Brosseron F, Teunissen C, Zetterberg H, Jacobs AH et al: Neuroinflammation in Alzheimer disease. Nat Rev Immunol 2025, 25(5):321–352.

51. Hoefs SJ, Dieteren CE, Distelmaier F, Janssen RJ, Epplen A, Swarts HG, Forkink M, Rodenburg RJ, Nijtmans LG, Willems PH et al: NDUFA2 complex I mutation leads to Leigh disease. Am J Hum Genet 2008, 82(6):1306–1315.

52. Peralta S, Torraco A, Wenz T, Garcia S, Diaz F, Moraes CT: Partial complex I deficiency due to the CNS conditional ablation of Ndufa5 results in a mild chronic encephalopathy but no increase in oxidative damage. Hum Mol Genet 2014, 23(6):1399–1412.

53. Mayr JA, Havlickova V, Zimmermann F, Magler I, Kaplanova V, Jesina P, Pecinova A, Nuskova H, Koch J, Sperl W et al: Mitochondrial ATP synthase deficiency due to a mutation in the ATP5E gene for the F1 epsilon subunit. Hum Mol Genet 2010, 19(17):3430–3439.

54. Wells L, Yang Q, Iorio C, Peragerasingam M, Campbell Z, Ng AC-H, Reeks C, Yee S-P, Screaton RA: ROMO1 and Mitochondrial Complex II/SDH are Required for Spare Respiratory Capacity and Glucose Homeostasis in Mice. bioRxiv 2024:2021.2004.2028.441811.

55. Chen J, Cai M, Zhan C: Neuronal Regulation of Feeding and Energy Metabolism: A Focus on the Hypothalamus and Brainstem. Neurosci Bull 2025, 41(4):665–675.

56. Pervolaraki E, Hall SP, Foresteire D, Saito T, Saido TC, Whittington MA, Lever C, Dachtler J: Insoluble Abeta overexpression in an App knock-in mouse model alters microstructure and gamma oscillations in the prefrontal cortex, affecting anxiety-related behaviours. Dis Model Mech 2019, 12(9):dmm040550.

57. Whyte LS, Hemsley KM, Lau AA, Hassiotis S, Saito T, Saido TC, Hopwood JJ, Sargeant TJ: Reduction in open field activity in the absence of memory deficits in the App(NL-G-F) knock-in mouse model of Alzheimer’s disease. Behav Brain Res 2018, 336:177–181.

58. Ulluwishewa D, Anderson RC, McNabb WC, Moughan PJ, Wells JM, Roy NC: Regulation of Tight Junction Permeability by Intestinal Bacteria and Dietary Components1,2. The Journal of Nutrition 2011, 141(5):769–776.

59. Zhu Y, Chen B, Zhang X, Akbar MT, Wu T, Zhang Y, Zhi L, Shen Q: Exploration of the Muribaculaceae Family in the Gut Microbiota: Diversity, Metabolism, and Function. Nutrients 2024, 16(16):2660.

60. Prasoodanan P. K V, Sharma AK, Mahajan S, Dhakan DB, Maji A, Scaria J, Sharma VK: Western and non-western gut microbiomes reveal new roles of Prevotella in carbohydrate metabolism and mouth–gut axis. npj Biofilms and Microbiomes 2021, 7(1):77.

61. Douros A, Ante Z, Fallone CA, Azoulay L, Renoux C, Suissa S, Brassard P: Clinically apparent Helicobacter pylori infection and the risk of incident Alzheimer’s disease: A population-based nested case-control study. Alzheimers Dement 2024, 20(3):1716–1724.

62. Jimenez-Garcia AM, Villarino M, Arias N: A systematic review and meta-analysis of basal microbiota and cognitive function in Alzheimer’s disease: A potential target for treatment or a contributor to disease progression? Alzheimers Dement (Amst) 2024, 16(4):e70057.

63. Hao W, Liu Y, Liu S, Walter S, Grimm MO, Kiliaan AJ, Penke B, Hartmann T, Rube CE, Menger MD et al: Myeloid differentiation factor 88-deficient bone marrow cells improve Alzheimer’s disease-related symptoms and pathology. Brain 2011, 134(Pt 1):278–292.

64. Luo Q, Schnoder L, Hao W, Litzenburger K, Decker Y, Tomic I, Menger MD, Liu Y, Fassbender K: p38alpha-MAPK-deficient myeloid cells ameliorate symptoms and pathology of APP-transgenic Alzheimer’s disease mice. Aging Cell 2022, 21(8):e13679.

65. Parker A, Fonseca S, Carding SR: Gut microbes and metabolites as modulators of blood-brain barrier integrity and brain health. Gut Microbes 2020, 11(2):135–157.

66. Zhang L, Zhang S, Maezawa I, Trushin S, Minhas P, Pinto M, Jin LW, Prasain K, Nguyen TD, Yamazaki Y et al: Modulation of mitochondrial complex I activity averts cognitive decline in multiple animal models of familial Alzheimer’s Disease. EBioMedicine 2015, 2(4):294–305.

67. Ferretti L, McCurry SM, Logsdon R, Gibbons L, Teri L: Anxiety and Alzheimer’s disease. J Geriatr Psychiatry Neurol 2001, 14(1):52–58.

68. Teri L, Ferretti LE, Gibbons LE, Logsdon RG, McCurry SM, Kukull WA, McCormick WC, Bowen JD, Larson EB: Anxiety of Alzheimer’s disease: prevalence, and comorbidity. J Gerontol A Biol Sci Med Sci 1999, 54(7):M348–352.

69. Palmer K, Berger AK, Monastero R, Winblad B, Backman L, Fratiglioni L: Predictors of progression from mild cognitive impairment to Alzheimer disease. Neurology 2007, 68(19):1596–1602.

70. Ko YK, Kim E, Lee EJ, Nam SJ, Kim Y, Kim S, Choi SY, Kim HY, Choi Y: Enrichment of infection-associated bacteria in the low biomass brain bacteriota of Alzheimer’s disease patients. PLoS One 2024, 19(2):e0296307.

71. Salzberg SL, Breitwieser FP, Kumar A, Hao H, Burger P, Rodriguez FJ, Lim M, Quinones-Hinojosa A, Gallia GL, Tornheim JA et al: Next-generation sequencing in neuropathologic diagnosis of infections of the nervous system. Neurol Neuroimmunol Neuroinflamm 2016, 3(4):e251.

72. Mone Y, Earl JP, Krol JE, Ahmed A, Sen B, Ehrlich GD, Lapides JR: Evidence supportive of a bacterial component in the etiology for Alzheimer’s disease and for a temporal-spatial development of a pathogenic microbiome in the brain. Front Cell Infect Microbiol 2023, 13:1123228.

73. Beydoun MA, Beydoun HA, Hossain S, El-Hajj ZW, Weiss J, Zonderman AB: Clinical and Bacterial Markers of Periodontitis and Their Association with Incident All-Cause and Alzheimer’s Disease Dementia in a Large National Survey. J Alzheimers Dis 2020, 75(1):157–172.

74. Dominy SS, Lynch C, Ermini F, Benedyk M, Marczyk A, Konradi A, Nguyen M, Haditsch U, Raha D, Griffin C et al: Porphyromonas gingivalis in Alzheimer’s disease brains: Evidence for disease causation and treatment with small-molecule inhibitors. Sci Adv 2019, 5(1):eaau3333.

