## Supplemental Figure 1 for "Gut miRNA regulates gut microbiome and Alzheimer’s pathology in *App*-knock-in mice"

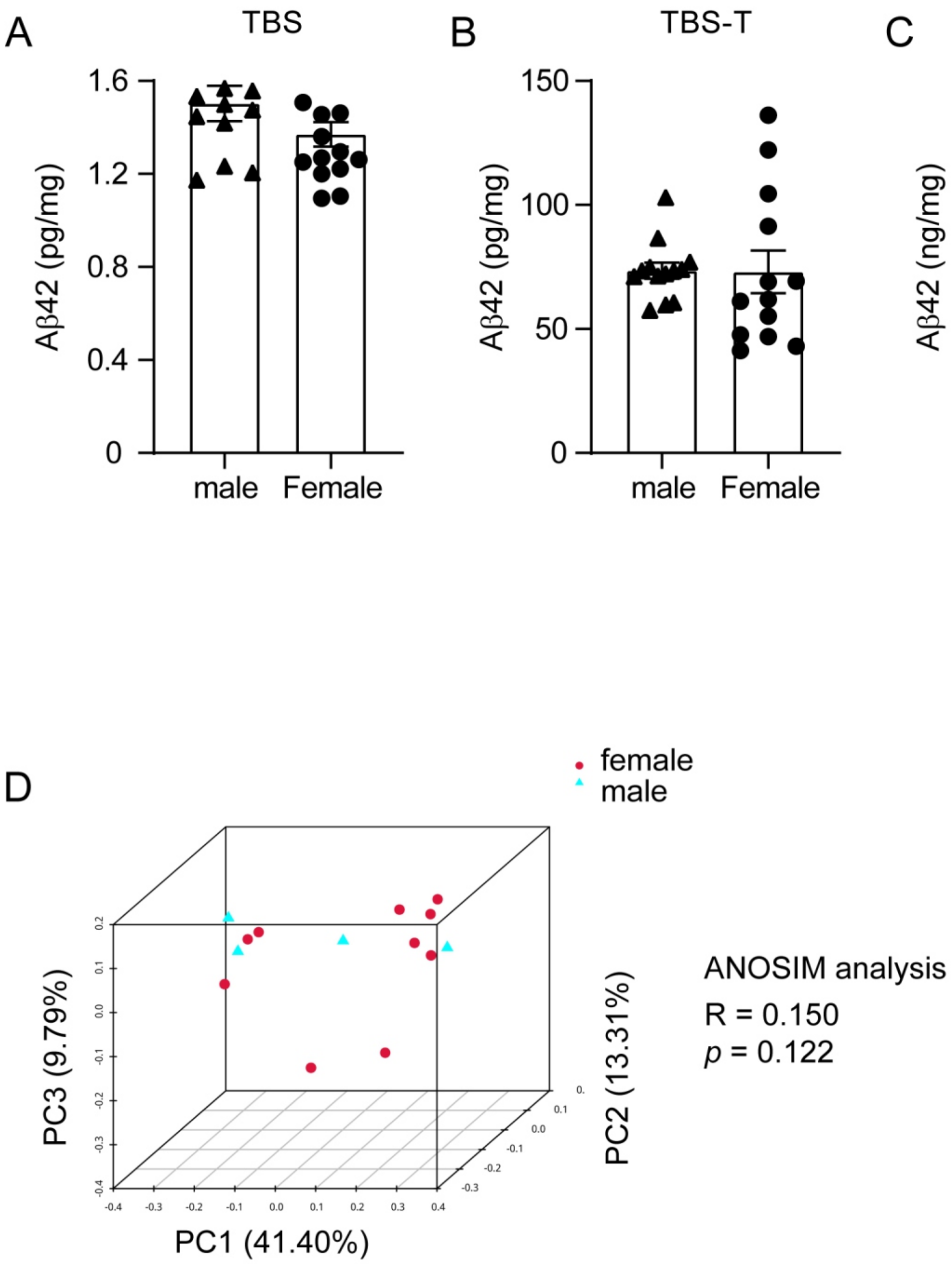


Supplementary Figure 1, Sex does not affect Aβ42 level in the brain and bacterial composition in the gut. A – C: Cortex and hippocampus tissue was collected from 9-month-old male and female App(ki/ki) mice, homogenized in TBS buffer plus 1% Triton-100 and analyzed for Aβ42 levels using an ELISA kit. Aβ42 levels were normalized by the sample's protein concentration, which was determined by Bradford assay. Sex had no significant effect on cerebral Aβ42 concentrations in App(ki/ki) mice.
t test, n = 12 and 15, for male and female mice, respectively.
D: Bacterial DNA was isolated from the cecum content of App(ki/ki) mice with and without deletion of dicer1 in gut epithelial cells and sequenced for the V3-V4 region of 16S rDNA. Principal coordinate analysis (PCoA) was used for β-diversity analysis of bacterial composition at the genus level. ANOSIM analysis showed that sex does not alter the structure of gut bacterial community.
n = 9 (male: 2) and 5 (male: 2) for AD mice with and without deletion of dicer1, respectively.
