## Supplemental Figure 2 for "Gut miRNA regulates gut microbiome and Alzheimer’s pathology in *App*-knock-in mice"

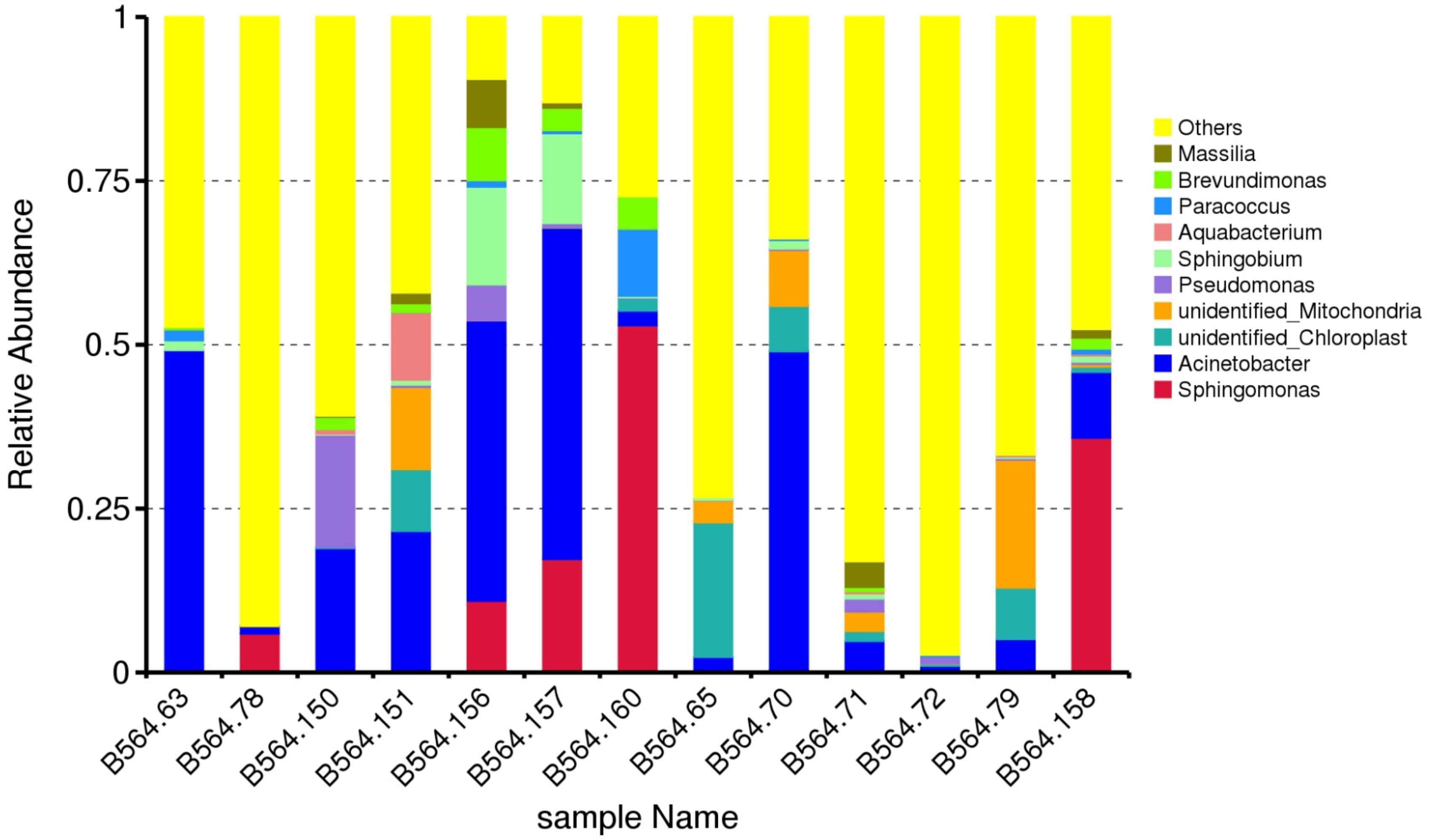


Supplementary Figure 2, Relative abundance of different bacterial genera in the brain tissues. Total DNA was isolated from brain tissues of App(ki/ki) mice with and without deletion of dicer1 in gut epithelial cells and sequenced for the V3-V4 region of 16S rDNA.
