## Supplemental Figure 3 for "Gut miRNA regulates gut microbiome and Alzheimer’s pathology in *App*-knock-in mice"

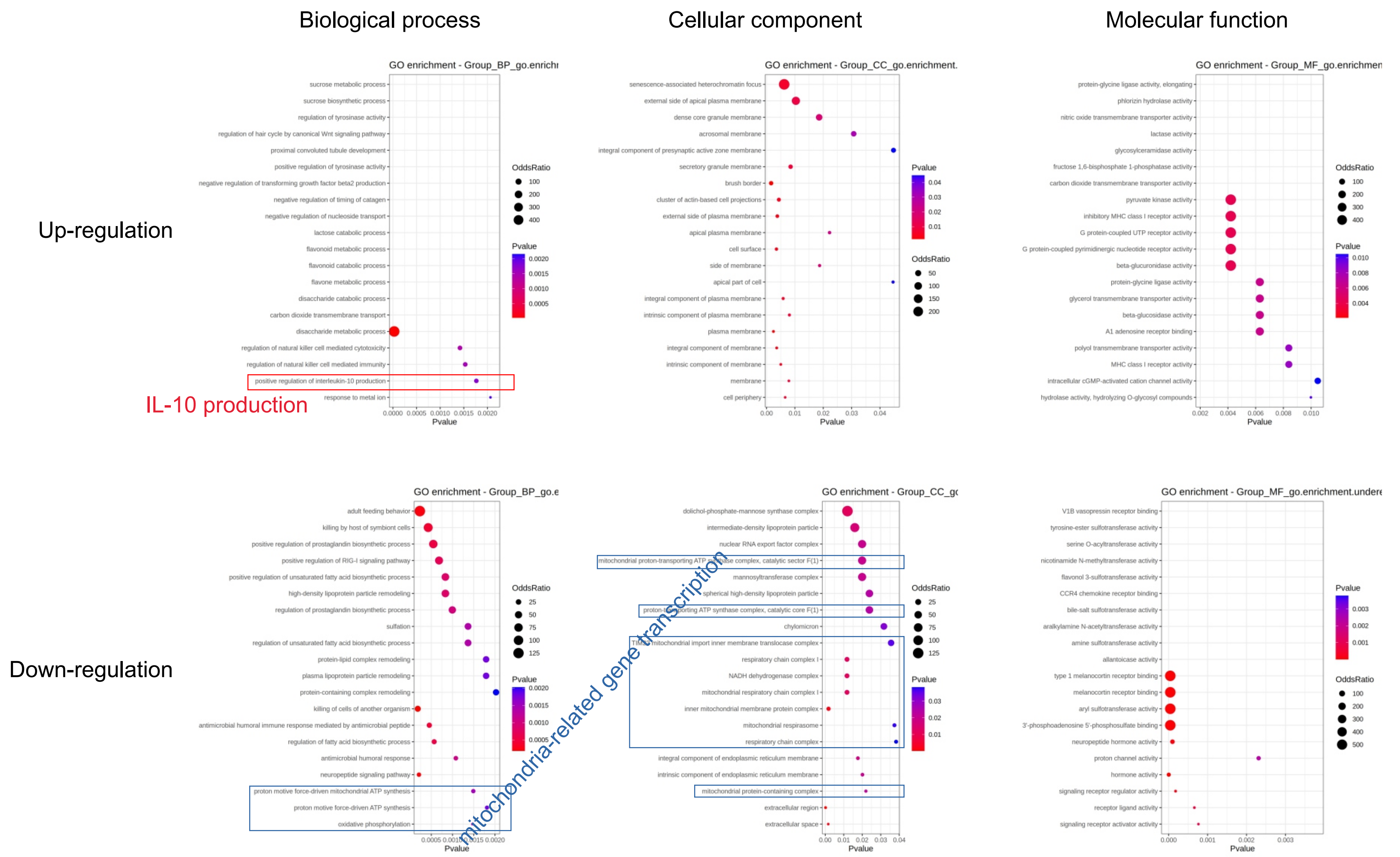


Supplementary Figure 3, GO enrichment analysis of differentially expressed genes.
Total RNA was isolated from brain tissue of App(ki/ki) mice with and without deletion of dicer1 in intestinal epithelial cells and sequenced for mRNA. Differentially expressed genes (DEGs) were identified with a *p*-value (instead of adjusted *p*-value) cut-off at 0.05 and a minimum absolute log2fold change 1. GO enrichment analysis showed that these DEGs with upregulated transcription are linked to IL-10 production in dicer1-deficient compared to dicer1 wild-type App(ki/ki) mice, whereas the DEGs with downregulated transcription are enriched in mitochondria-related genes.
n = 7 and 12 for mice with and without deletion of dicer1, respectively.
